# Prolactin Shapes Cortical Plasticity in Fathers

**DOI:** 10.1101/2025.10.20.683400

**Authors:** B. Haimson, T. Preminger, P.R. Rayi, H. Sagi, A. Binshtok, A. Mizrahi

**Author notes:** Corresponding author: Adi Mizrahi.

## Abstract

Caregiving alters the mammalian brain to support infant survival. While hypothalamic circuits are known to drive innate parental behaviors, information about the effect of parenthood on sensory processing and perception remain scarce, especially in fathers. We longitudinally imaged the same neurons in male mouse primary auditory cortex (ACx) before, during, and after fatherhood using two-photon calcium imaging. Behaviorally, males shifted from pup-directed aggression and self-oriented behaviors to transient parental care. In ACx, fathers showed enhanced and faster discrimination of pup ultrasonic vocalizations compared with matched narrowband noise. Slice electrophysiology revealed increased intrinsic excitability accompanies fatherhood. Mechanistically, we found elevated prolactin signaling in ACx, and manipulating prolactin levels modified the improvement in neuronal discriminability. Together, these results identify–in males–cortical plasticity and link prolactin to experience-dependent plasticity of sensory circuits that support caregiving. Our findings outline a father-specific mechanism that differs in key aspects from similar mechanisms in mothers.

## Introduction

Newborn animals’ complete dependence on parental care is a key selective pressure that has shaped the evolution of diverse caregiving strategies across species ^1,2^. To support efficient caregiving, parents undergo widespread neural adaptations across multiple brain regions. Considerable research on parental plasticity has focused on hypothalamic circuits, particularly those modulated by oxytocin and estrogen ^3–8^, revealing innate mechanisms of parental care. Yet, it has long been clear that there is a significant experience-dependent component to parental behavior, and there is growing evidence that parenthood also modulates early sensory processing, especially in mothers^9–15^. Despite progress in our understanding of maternal plasticity, we know far less about how these circuits contribute to parenting, and what are their underlying mechanisms.

In rodents, females undergo profound behavioral and neural changes during motherhood. Sensory regions, including auditory cortex and olfactory bulb, exhibit changes that enhance the detection and interpretation of offspring-derived cues ^9–13,15–27^. For example, compared to naïve females, mothers show rapid and efficient responses to pup signals, such as ultrasonic vocalizations (USVs), which serve as key components in maternal caregiving behaviors ^24,28^. In contrast, while males also exhibit striking behavioral transitions upon becoming fathers, shifting from aggression or infanticide towards pups to active nurturing ^29–31^, the corresponding neural adaptations remain poorly understood. Whether fatherhood induces plasticity in sensory regions is not known.

A major difference between mothers and fathers is the distinct hormonal milieu, both overall and specifically in the hormones that shape parental behavior. Maternal plasticity is tightly linked to peripartum hormonal shifts, especially estrogen and oxytocin ^4,24,32–41^. In males, however, these hormones play little to no role in fatherhood. Instead, recent findings implicate prolactin, a hormone traditionally associated with changes in maternal physiology like lactation, as a key modulator of paternal behavior ^42,43^. However, these studies focused only on prolactin-related modulation of hypothalamic networks. For example, prolactin receptor gene knockout in the medial preoptic area significantly reduced parental behavior in male mice ^43^. Information about the involvement of prolactin in fatherhood-related plasticity in other brain regions is still lacking.

In this study, we examined neuronal plasticity in the primary ACx of male mice across the transition to fatherhood. We focused on the cortical representation of pup USVs as a salient social cue and found that USVs are encoded differently in active fathers. We further identified increased prolactin signaling as a modulator of these cortical responses. Together, our experiments reveal fatherhood-associated changes in cortical physiology and provide new insight into male-specific mechanisms of paternal brain plasticity.

## Results

### Male mice transition into parental care when they become fathers

To study parental plasticity associated with fatherhood, we first characterized its behavioral manifestations. We used the ICR:CD1 mouse model, which is known to exhibit pronounced parental behaviors ^28,44^. Male mice were housed with a female, and remained with her following mating, through parturition, until their own pups were at the age of 14-21 days.

Since mating alone can induce behavioral and hormonal changes ^42,45–47^, we started by assessing parental behaviors in males 3-5 days post-mating (‘Post mating males’), and then again 4-6 days after parturition (‘Fathers’; **Fig. 1a**). We evaluated paternal behaviors through 15-minute pup interaction assays, in which four pups were placed on the opposite side of the home cage to be retrieved (**Fig. 1b**).

**Figure 1.**
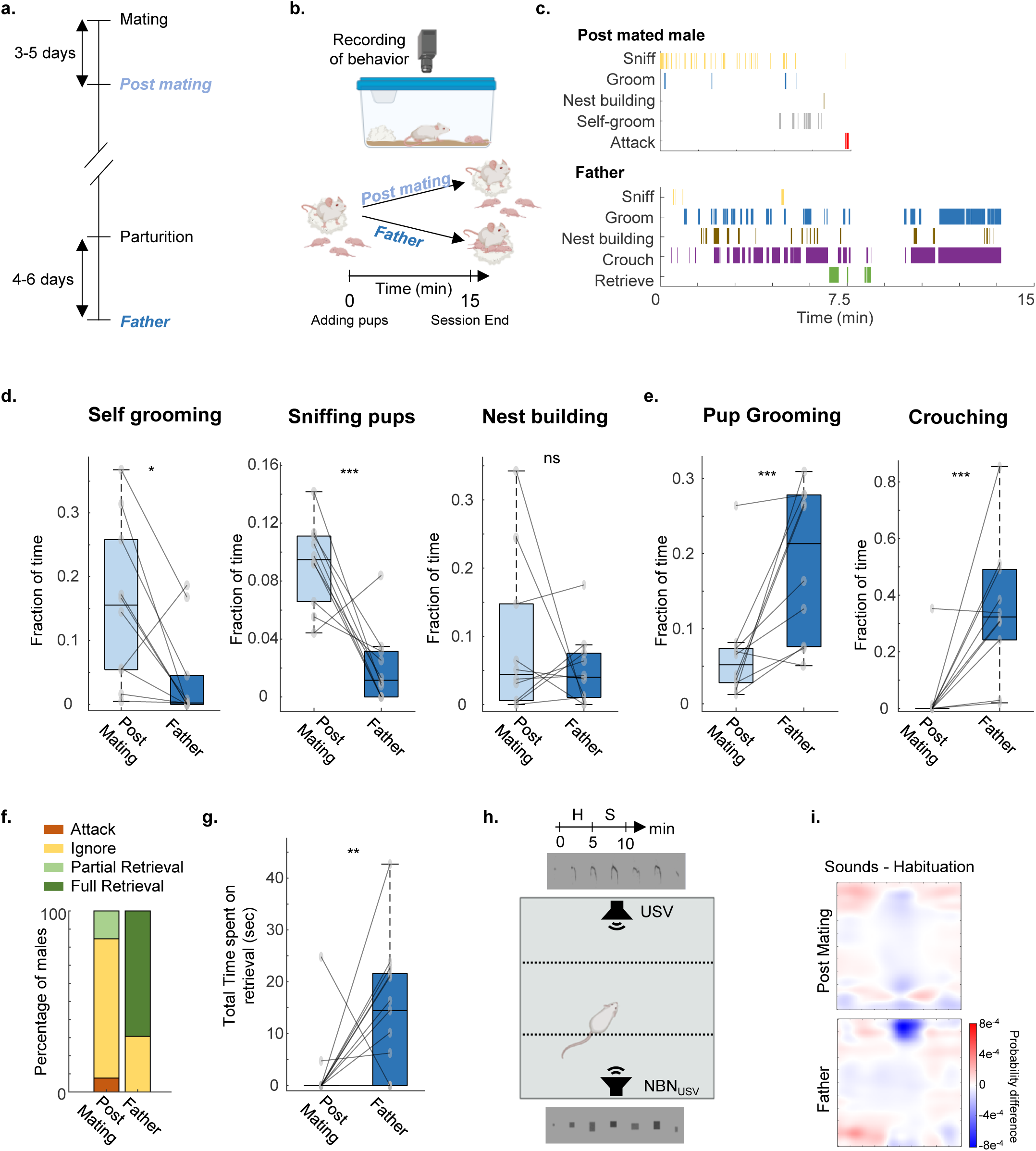
Fatherhood induces a transient shift to paternal care. (a) Experimental timeline depicting the two assessment time points: Post-mating (3– 5 days after mating) and Father (4–6 days after parturition). (b) Schematic illustration of the behavioral tracking setup, monitoring mouse actions over the 15-min pup retrieval assay in the home cage. (c) Representative behavioral ethogram of one representative mouse, showing the transition from aggressive and self-centered behavior before fatherhood to parental care as a father. (d-e) Fraction of time dedicated for self-grooming, pup sniffing and nest building (d) and pup grooming and crouching (e) in the post-mating stage and as fathers (n = 10 mice). (f) Proportion of mice exhibiting pup retrieval behavior as post mating males and as fathers (n = 18 mice). (g) Total time spent on retrieving pups behavior as post mating males and as fathers (n = 10 mice). (h) Schematic illustration of the test timeline (top) and the experimental setup of an auditory spatial-preference test with ultrasonic vocalizations (USVs) and narrowband noise (NBN) speakers located in opposite corners of an open arena. H: Habituation, S: Sounds ON. (i) Heatmap of spatial preference (calculated as the spatial preference. Location during sounds ON subtracted by the period of habituation, average of all mice). Top: post mating males; Bottom: fathers (n = 13 mice). Data are median ± IQR; comparisons by Wilcoxon signed-rank test; ns p > 0.05, *p < 0.05, **p < 0.01, ***p < 0.005.

Males exhibited aggressive or self-focused behaviors (i.e., self-grooming) during the ‘post-mating’ period, which transitioned into active parental care once they became fathers (**Fig. 1c–e**, **Extended Data Fig. 1**). Most post-mated males ignored pups, with only a few mice exhibiting delayed or partial retrieval. As fathers, however, the majority of male mice rapidly retrieved all pups (**Fig. 1f,g, Extended Data Fig. 2d–f**).

In a subset of males, we tested the transient nature of these behaviors. Specifically, we tested fathers that were isolated from the mother and pups for 7 days (a group we named ‘Post-Fatherhood Males’). Post-fatherhood males showed a loss of most paternal behaviors, towards new young pups (P3-5) (**Extended Data Fig. 2**). The loss of parental behaviors after separation suggests that active fatherhood depends on ongoing social and sensory experience with the pups.

### Valence of pup vocalizations during fatherhood becomes aversive

Parental behavior is multifaceted, involving numerous brain regions and sensory modalities ^12,48–50^. Here, we focused on the auditory domain, particularly in light of the novel acoustic environment that mice encounter upon transition to parenthood. Specifically, pups emit USVs that are a critical component of effective parent–pup interactions ^21,28^. Moreover, the ACx is known as essential for maternal behavior and undergoes significant plasticity in mothers, particularly in how cortical neurons respond to pup USVs ^11–16,24,28^.

To test whether fathers account for pup USV differently, we performed an auditory place preference test (**Fig. 1h**). We compared the spatial preference of male mice at different parental stages towards pup USV versus a control stimulus of corresponding narrowband noise (NBN) with the same sound energy and temporal structure as the USV but lacking amplitude and frequency modulations. We presented the two sounds from different directions in an open field arena and let mice 5 minutes to freely explore it (**Fig. 1h; Extended Data Fig. 3a**). Post-mated males showed no spatial preference for either sound, indicating that prior to active fatherhood the pup USV lacks intrinsic valence. However, fathers showed avoidance of the pup-USV zone that emerged in the last 1–2 minutes of the 5-min assay (**Fig. 1i, Extended Data Fig. 3b-c**); a result which is consistent with previous findings in dams, indicating that fathers too perceive USV as aversive stimuli ^51,52^. Importantly, total locomotor activity remained stable across conditions (**Extended Data Fig. 3d, e**), confirming that avoidance behavior was not due to reduced exploratory drive. Thus, the transition to fatherhood is accompanied by a shift in auditory valence of USV from neutral to aversive.

### Fatherhood transiently alters pure tone tuning and responses to pup USVs

To date, data on fatherhood-induced cortical plasticity remains scarce ^53,54^. To investigate this, we performed longitudinal *in vivo* two-photon calcium imaging in awake mice that were injected with AAV9-hsyn-GCaMP6s-P2A-nls-dTomato into the ACx. We measured neuronal responses to various sounds in male mice at three time points: ‘PM’: post-mating (before fatherhood), ‘F’: fathers (5 days postpartum), and ‘PF’: post-fatherhood (7 days after separation from the dam and pups). We performed imaging in head-restrained, awake mice, under passive listening conditions (**Fig. 2a-c**; N= 856, 861, and 594 neurons for each time point, respectively (PM & F, n=9 mice; PF, n=6 mice).

**Figure 2.**
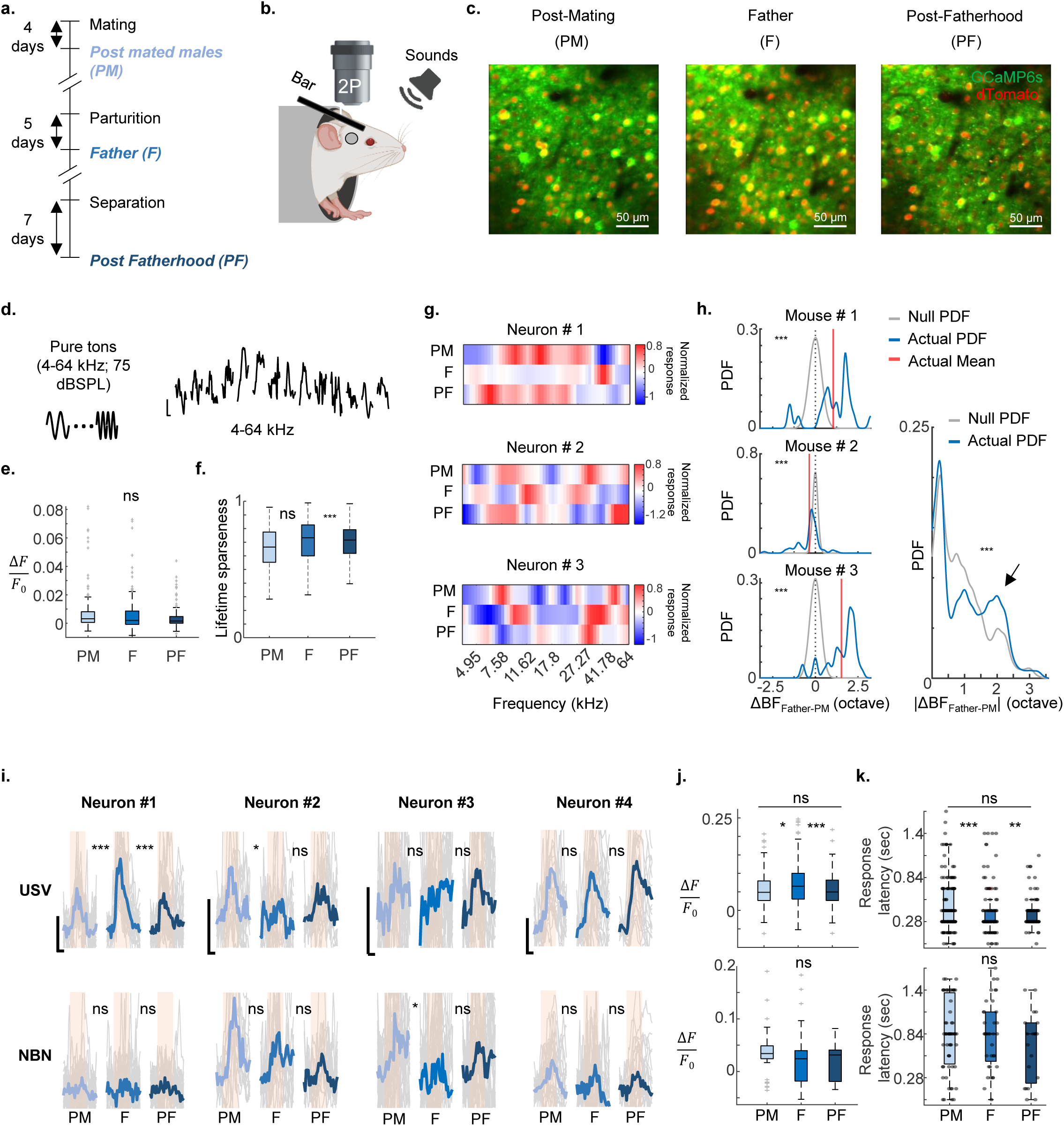
Fatherhood induces transient shifts in frequency tuning and responses to pup calls in primary auditory cortex. (a) Experimental timeline depicting the three assessment time points: Post-mating (3–5 days after mating), Father (4–6 days after parturition) and Post fatherhood (7 days after separation from dam and pups). (b) Experimental setup – two-photon imaging from an awake mouse passively listening to sounds. (c) Representative in vivo two photon micrographs from one mouse at the different time points. Green, GCaMP6s; red, dTomato. (d) Left: Single neuron tuning curves were constructed by playing pure tones from 4 to 64 kHz logarithmically spaced at 75dBSPL. Right: example tuning of a single neuron. (e) Population ΔF/F₀ average responses to all pure tones at the three time points (n = 280, 299 and 157 responsive neurons, from 5,5 and 3 mice, at post mating, fatherhood and post fatherhood stages, respectively). LME followed by FDR for multi comparison correction. (f) Lifetime sparseness at the three time points (n = 280, 299 and 157 responsive neurons, from 5,5 and 3 mice, at post mating, fatherhood and post fatherhood stages, respectively). P-values: PM vs. F: 0.07. PM vs. PF: 0.1. F vs. PF: 0.0022. LME followed by FDR for multi comparison correction. (g) Tuning curves of three example neurons, at three different time points. Note the transient changes in neurons #1 and #2, but not in neuron #3. (h) Probability distribution function (PDF) of the change in neuronal best frequency (BF) between the post mating and father time points. The null distribution assuming random changes is indicated in gray (dotted line, mean). Mean of the actual distribution (blue) is indicated in red. Left: PDF’s for three single mice (p-values: 0.0018, 3.67e-6 and 4.1e-7, for the three mice, respectively, Wilcoxon signed rank test – compared to distribution with median zero). Right: the absolute change in BF compared to the null distribution for all mice (n = 295 neuron from 3 mice, Permutation test p-value (10000 shuffles) < 0.0001). The black arrow indicates an increase in BF change which does not appear in the null distribution. X-axis values are in octaves. (i) Calcium traces from four representative neurons in response to USVs (upper row) and NBN (lower row). Each neuron is shown at the three time points (PM, F, PF). Mean ΔF/F₀ amplitudes are shown in a thick line; single trials in grey. The pink shade indicates time of stimulus exposure. Scale bar: 0.04 ΔF/F₀ /1 sec. (j) Mean ΔF/F₀ amplitudes in response to USVs (top) and NBN (top) at the three time points (n = 165, 140 and 105 and n = 78, 80 and 40 USVs or NBN responsive neurons for 3 time points, respectively from 9 mice). P-values (USVs): PM vs. F: 0.043. PM vs. PF: > 0.05. F vs. PF: 0.0003. LME followed by FDR for multi comparison correction. (k) Latency of responses to USVs (top) and NBN (bottom) at the three time points (n = 165, 140 and 105 and n = 78, 80 and 40 USVs or NBN responsive neurons for 3 time points, respectively from 9 mice). P-values (USVs): PM vs. F: 6.17e-6. PM vs. PF: 0.14. F vs. PF: 0.006. For NBN all p-values > 0.05. LME followed by FDR for multi comparison correction. Data are median ± IQR; comparisons by LME followed by FDR for multi comparison correction.

We started by measuring single-neuron tuning to evaluate basic auditory responses to pure tones **(Fig. 2d**; 4 to 64 kHz). Overall, we found no fatherhood-related change in mean evoked response or in lifetime sparseness (**Fig. 2e, f,** N= 470, 490, and 314 neurons for each time point, respectively (PM & F, n=5 mice; PF, n=3 mice). However, we found that individual cells exhibited transient shifts in best frequency (BF) during fatherhood **(Fig. 2g)**. Closer inspection of the distribution of BFs of all neurons showed a significant shift in tuning properties at the population level (**Extended Data Fig. 4a**). The observed distribution of BF changes was significantly higher than expected by chance **(Fig. 2h** and **Extended Data Fig. 4b**).

Notably, some neurons transiently shifted toward higher frequencies and others toward lower frequencies. This bidirectional change in tuning was also higher than chance (skewness = 0.54 vs. 0.86, for real data compared to chance, respectively, with individual neuron change up to 3.25 octaves), indicating genuine redistribution of stimulus-specific responses in ACx neurons during fatherhood (**Fig. 2h**).

Next, we measured neuronal responses to USV and their corresponding NBN. Single-neuron responses varied considerably across time, yet a considerable proportion of the neurons (22%) showed selective changes (increases or decreases) specifically during fatherhood (**Fig. 2i**). On average, response amplitudes to USV slightly increased (by ∼23% on average) following fatherhood, without similar effect to their corresponding NBN (**Fig. 2j**). This increase was not due to an increase in the fraction of the responsive neurons in the network (**Extended Data Fig. 4c, d**). Additionally, we found that response latency was significantly shorter in fathers and partially restored after fatherhood (**Fig. 2k**). This effect was observed only in responses to USV, but not to their corresponding NBN (**Fig. 2k**).

### Fatherhood is accompanied by better and faster cortical discriminability of pup USVs

Next, we asked if fathers exhibit improved pup vocalization coding. To address this, we assessed how neurons discriminated between USV and their corresponding NBN. To do so, we calculated a receiver operating characteristics curve between the distributions of each neuron’s responses to each sound and calculated its area under the curve (AUC). To compensate for the neuronal preferences of either sound, we used a modified version of the raw AUC value, indicated by a discriminability index, DI (see Methods).

Single-neuron discriminability changed in a heterogenous manner (**Fig. 3a**). On average, discriminability increased significantly during fatherhood and returned to baseline levels in post fatherhood (**Fig. 3b**). Neuronal discriminability was not only stronger, but also faster. Neurons in fathers discriminated USV from NBN, on average, 295 ms faster than in non-fathers (**Fig. 3c,d**). This effect did not stem from difference in the number of neurons that did not pass the ‘discriminability threshold’ (**Fig. 3e**, see Methods). Notably, these two effects––improvement in single-neuron discriminability and speed––were strongly correlated, suggesting that the same neuronal subpopulation contributes to both effects (**Fig. 3f, Extended Data Fig. 5a**).

**Figure 3.**
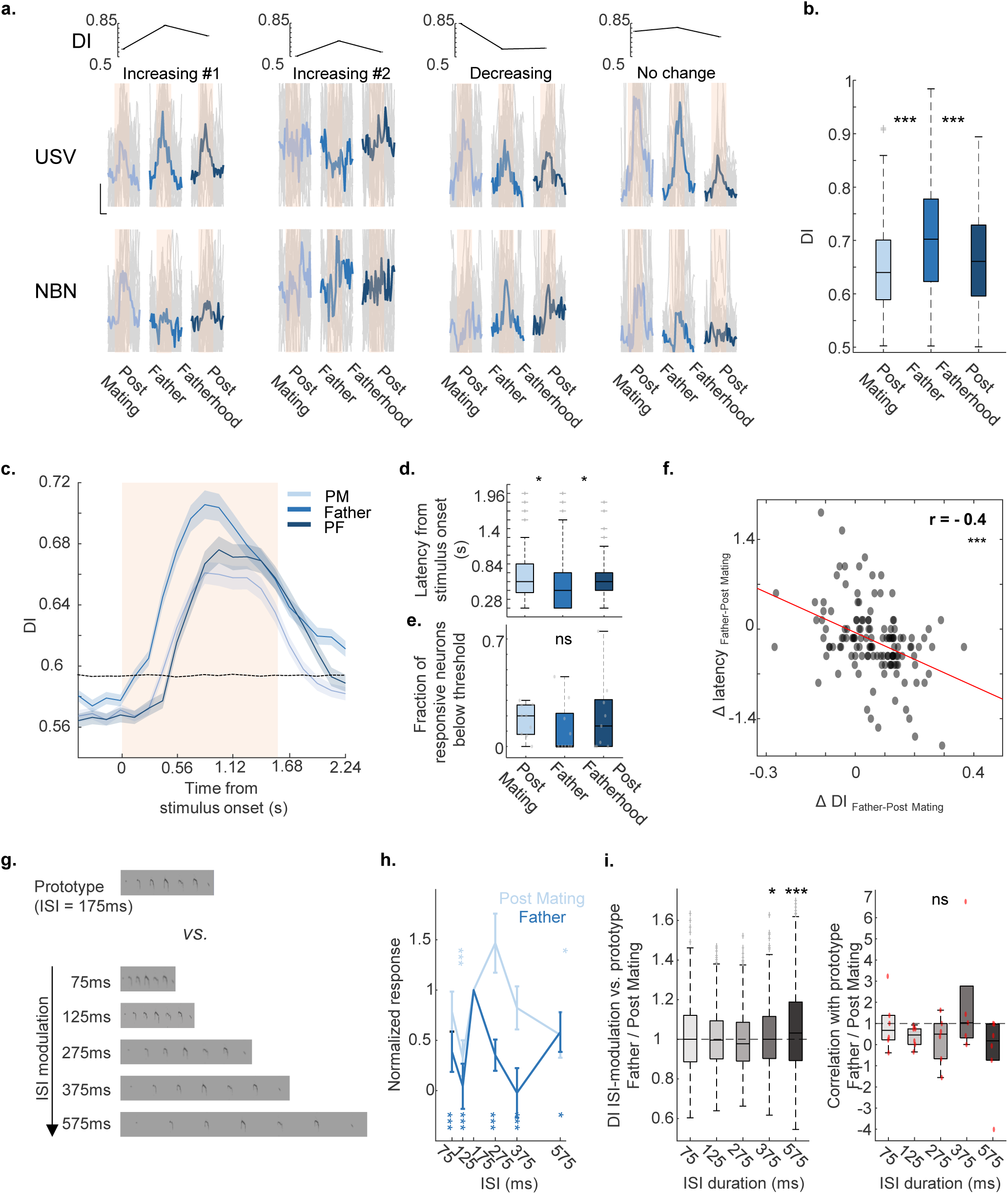
ACx neurons in fathers exhibit higher and faster discrimination of pup USVs. (a) Example calcium traces (ΔF/F) for both USVs and NBN from four representative neurons across the three time points of assessment. Upper row: the corresponding discrimination index (DI) for each of the time points. (b) Single-neuron DI for USV vs. NBN significantly increased during fatherhood (n= 191, 199 and 132 neurons (responsive to USVs or their corresponding NBN), from 9, 9 and 6 mice for the different time points). P-values: PM vs. F: 7.75e-10. F vs. PF: 5.9e-7. PM vs. PF: 0.42. LME followed by FDR for multi comparison correction. (c–d) Time-resolved decoding analysis for the different time points (c). There is significantly faster discrimination of USVs (vs NBNs) during fatherhood (n= 191, 199 and 132 neurons from 9, 9 and 6 mice). All neurons in this analysis were responsive to USVs or their corresponding NBN. P-values: PM vs. F: 0.013. F vs. PF: 0.036. LME followed by FDR for multi comparison correction. P-values: PM vs. F: 0.0074. F vs. PF: 0.0018. PM vs. PF: 0.42. LME followed by FDR for multi comparison correction. (e) The fraction of responsive neuron below threshold at each time point (n= 9 mice). (f) A correlation between the change in discriminability and change in discriminability latency (n= 219 neurons, from 6 mice). P-values: 1.72e-6, Pearson correlation. (g–i) Fathers showed improved discriminability for USV at varying ISIs. (g) spectrograms of USVs with different ISIs. (H). average response to temporally manipulated USVs in post mating males and in fathers. (I) left: the ratio between single neuron DI (prototype USV vs. manipulated versions) in fathers and the same DI as post mating males. Ratio above one indicates better discrimination in fathers, and below one, better discrimination at post mating stage. Right: similar ratio between the Pearson correlations of the whole population response. No significant generalization is evident in fathers. P-values h (from prototype): PM: 0.27, 1.392e-4, 0.11, 0.406, 0.0466. F: 0.0024, 2.5774e-5, 2.4172e-5, 3.91e-5, 0.0348, for 75,125,275,375 and 575 ms ISI, respectively (LME and FDR correction). P-values i (left): 375ms: 0.029, 575: 7.15e-8 (Paired t-test (different from 1), Bonferroni correction). Data are median ± IQR; comparisons by LME followed by FDR for multi comparison correction.

To test whether these changes were specific to USV, we examined responses to wriggling calls (WCs), another common type of pup call produced during competition for nursing ^55^. We compared single-neuron discriminability between WCs and their corresponding NBN. For WCs, neither the accuracy nor speed of discriminability differed (**Extended Data Fig. 5b-e**), suggesting that plasticity was selective towards USVs. These results also suggest that auditory experience alone (i.e., exposure to pup calls) is unlikely to be the only driver of this cortical plasticity to USVs in fathers.

The improved discriminability of single neurons we observed in fathers differs from previous findings in females, which showed generalization across acoustic variants of pup USV ^15,52^. To directly assess neuronal generalization in males, we adapted the paradigm from Schiavo et al. (2020) and presented temporally manipulated USVs with varying inter-syllable intervals (ISIs; **Fig. 3g**). Fathers showed no evidence of generalization; instead, they exhibited enhanced discriminability for temporally stretched calls at both the single-neuron and population levels (**Fig. 3h-i**). This phenotypic difference from mothers points to distinct information processing strategies in the ACx between males and females, which is likely mediated by distinct underlying neural mechanisms.

### Intrinsic excitability of ACx neurons is higher in fathers

We examined whether fatherhood affects the intrinsic excitability of auditory cortical neurons by performing whole-cell patch-clamp recordings from layer 2/3 pyramidal neurons in acute brain slices from post-mated males and fathers (**Fig. 4a**). Neurons from fathers exhibited significantly higher action potential (AP) firing rates in response to current injections compared to post-mated males (**Fig. 4b**). These neurons also show an increase in gain (**Fig. 4b**) and a decrease in the threshold current (rheobase, **Fig. 4c**). This increase in neuronal excitability was accompanied by elevated input resistance (**Fig. 4d**). Neurons from fathers also demonstrated a slight but significant depolarized resting membrane potential (**Fig. 4e**), an increase in AP threshold, a decrease in AP amplitude and maximal rate of AP rising (dV/dt_max_), without changes in AP half-width (**Extended Data Fig. 6**). Together, these data show that fatherhood is accompanied by alterations in the intrinsic properties of ACx neurons, leading to enhanced neuronal excitability in fathers.

**Figure 4.**
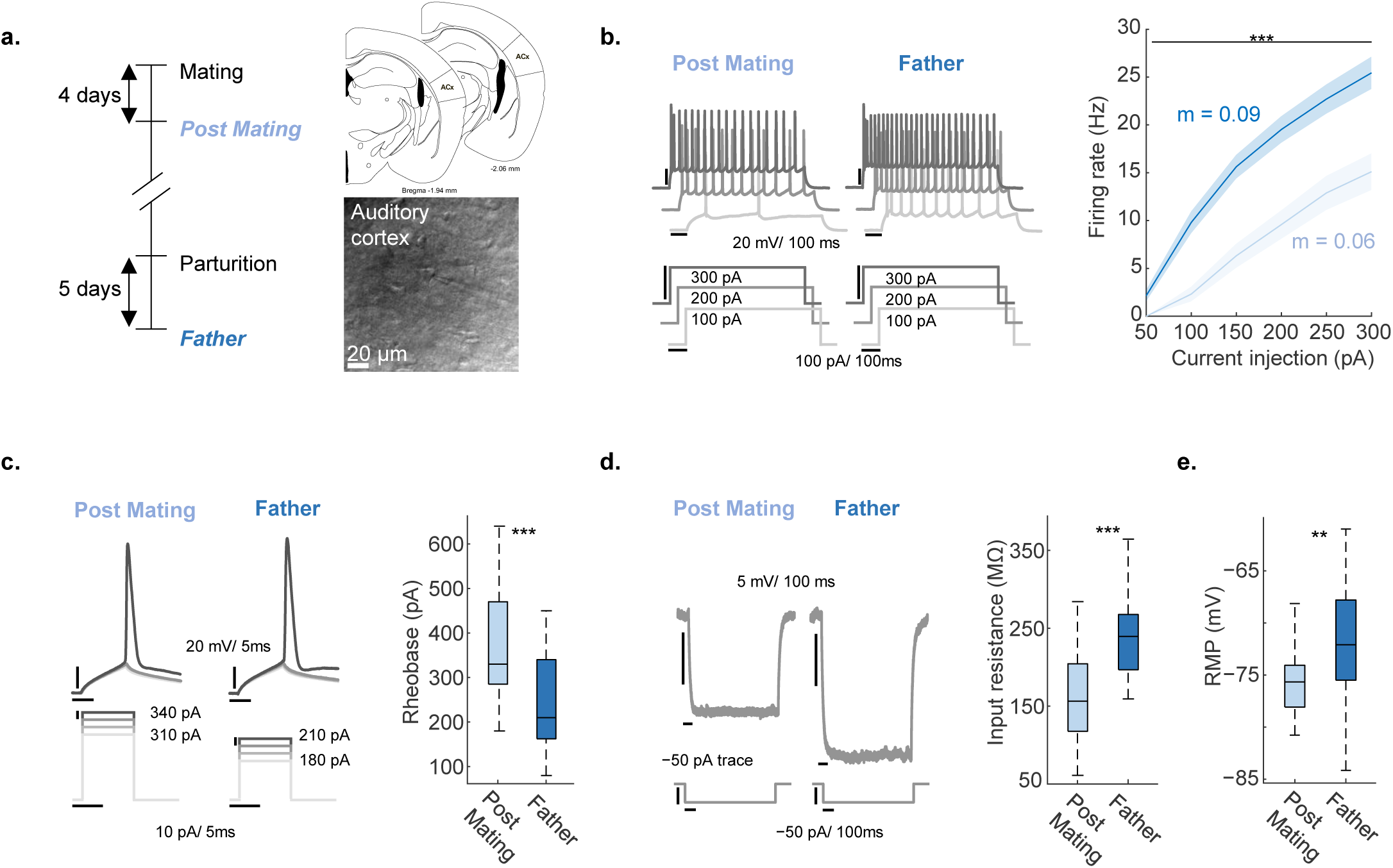
Intrinsic excitability of auditory cortical neurons is elevated in fathers. (a) Left: The two experimental groups used for slice electrophysiological recordings. Right (top): Schematic of the brain slices consisting ACx regions used in the experiment. Right (bottom): IR-DIC image of ACx pyramidal neuron with the recording pipette. (b) Left: Representative traces of action potential firing evoked by 1s depolarizing current steps (indicated below) from a neuron from a post mating male and a neuron from a father. Right: Firing rate versus current injections (*f*-I curves) in post-mated males and fathers (p-values: 0.0019, 0.0001, 0.0001, 0.0001, 0.0004, 0.0007, for 50, 100, 150, 200, 250, and 300 pA current injections, respectively. Mann–Whitney U test followed by FDR correction). The *f*-I slope of the fitted regression line for each stage is indicated by m in the corresponding color. (c) Representative traces of single action potentials from fathers and post-mating males evoked by 10 ms current steps. Right: threshold current (rheobase) is significantly lower in fathers (p-value: 4.3259e-04). (d) Left: Examples of changes in membrane voltage in response to application of hyperpolarizing current step of −50 pA. Right: Calculated input resistance was significantly higher in fathers (p-value: 9.1547e-05). (e) Resting membrane potential is depolarized in fathers (p-value: 0.0052). The data in this figure consist of n=19 and 27 neurons, obtained from 5 and 7 post mating and father mice, respectively (1-2 slices per mouse). Data are median ± IQR; *p<0.05, **p<0.01, **p<0.005 by unpaired t-test or Mann–Whitney U test.

### Prolactin signaling is stronger in the ACx of fathers

The transition to parenthood in mammals entails a series of hormonal shifts that promote both physiological changes and neural adaptations that are crucial for ensuring offspring survival ^56^. While hormones such as estrogen and oxytocin are classically associated with maternal behaviors ^32,57^, recent findings suggest that prolactin plays a prominent role in fatherhood ^42,43^. Based on those findings, we tested whether prolactin signaling also contributes to cortical plasticity in fathers.

We first measured levels of phosphorylated STAT5 (pSTAT5), a well-established downstream marker of prolactin receptor activation ^42,43,58,59^. Fathers exhibited a robust increase in pSTAT5 expression in the ACx (**Fig. 5a,b**), with the strongest signal observed in layers 2/3 and 4 (**Fig. 5c**). This increase was transient, returning to post-mating levels after the cessation of fatherhood (**Fig. 5a-c**). As a positive control, we confirmed the elevation of pSTAT5 in the medial preoptic area, a hypothalamic region known to be regulated by prolactin and critical for parental behavior in both sexes ^60^ (**Extended Data Fig. 7**). In contrast, and consistent with prior studies, we did not detect changes in the periaqueductal gray, which is known to exhibit decreased activity during fatherhood ^61^. Other sensory areas like the visual cortex showed no change, suggesting that prolactin signaling enhancement is not global (**Extended Data Fig. 7**).

**Figure 5.**
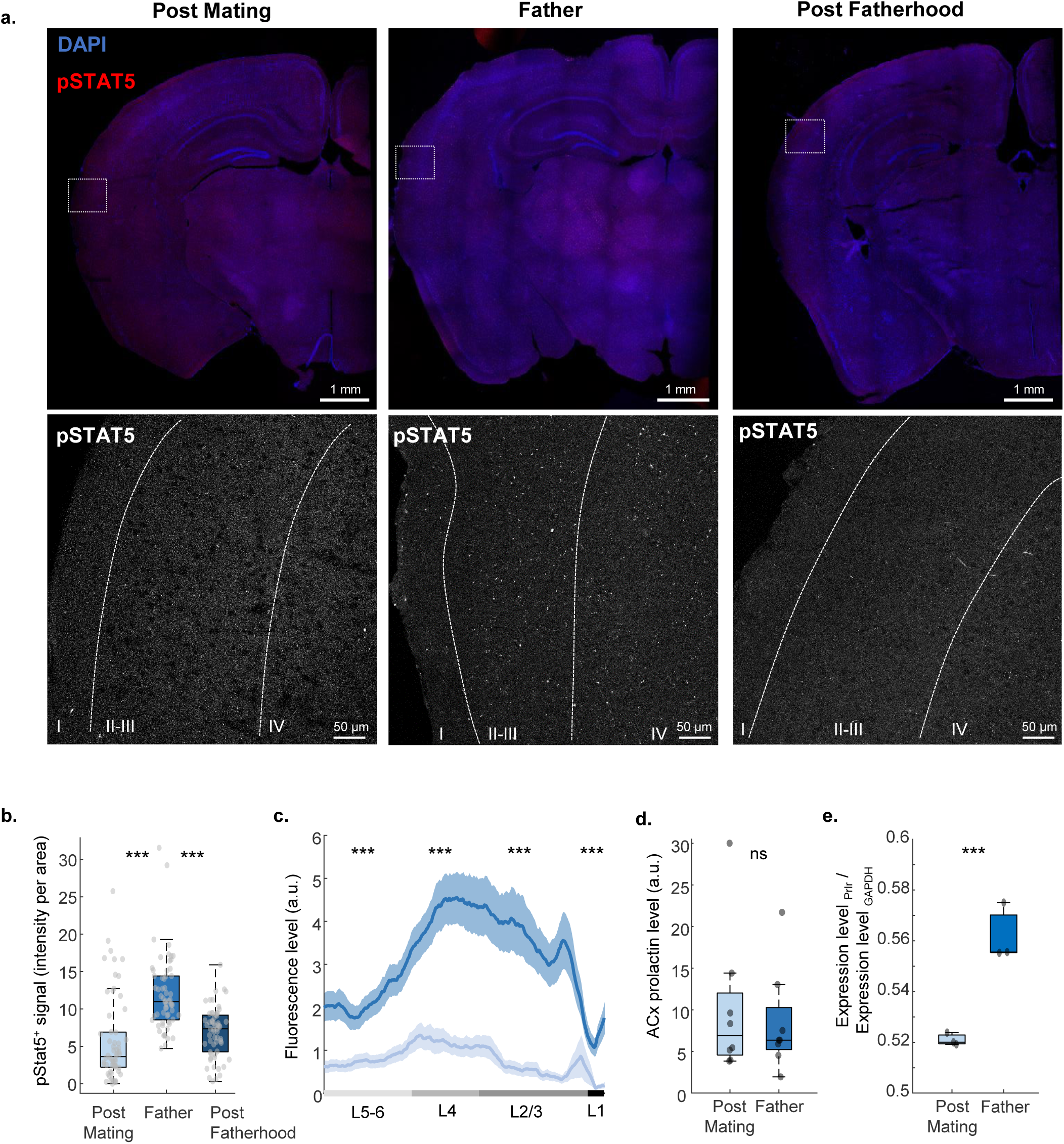
Prolactin signaling is higher in the auditory cortex of fathers. (a) Representative fluorescent micrographs of pSTAT5 immunostaining in ACx in three experimental groups. Top: low resolution coronal slice. Bottom: high magnification images of the ACx (marked with a white triangle in the top). (b–c) Quantification of pSTAT5+ signal in ACx across the three groups (b). Layer specific pSTAT5+ signal in ACx in post-mated males and fathers. p-values (b). One-way ANOVA: 2.876e-11. PM vs F: 9.6e-10, F vs PF: 2.7e-8, PM vs PF 0.42, after Tukey-Kramer multiple comparison correction. Data were collected from 12 brain slices per mouse, across 5 mice. LME corrected p values (c): L1:3.4e-21, L2-3: 7.9e-20, L4: 3.7e-18, L5-6: 8.7e-17. (d) Level of prolactin in postmated males and fathers (n = 8 and 8 mice, for both post-mating and father time points. p = 0.66, Unpaired t-test). (e) Relative expression levels of the long isoform of prolactin receptor (Prlr-L) in ACx of fathers (n = 3 and 3 mice, for both post-mating and father time points. p = 0.0028, Unpaired t-test). Data are median ± IQR; *p<0.05, **p<0.01, **p<0.005 by unpaired t-test or One-way ANOVA followed by Tukey-Kramer correction for multiple comparisons or LME followed by FDR for multi comparison correction.

The elevated pSTAT5 levels might stem from multiple sources, including an increase in circulating prolactin or changes in local receptor expression. To distinguish between these possibilities, we first measured prolactin levels in the ACx using enzyme-linked immunosorbent assay and found no significant differences between fathers and post-mated males (**Fig. 5d**). This result is consistent with prior findings from C57BL6 mice ^42^, and it suggests that the increased pSTAT5 signal is not driven by elevated levels of prolactin.

Because prolactin levels in the ACx did not differ between groups (**Fig. 5d**), we asked whether fathers’ ACx becomes more sensitive to prolactin. One parsimonious mechanism is upregulation of the prolactin receptor. In mouse, prolactin receptors include long and truncated short isoforms, which differ in their ability to propagate STAT5-dependent signaling. We quantified prolactin receptor long transcripts by qPCR and found a significant increase in fathers’ ACx (**Fig. 5e**). We interpret transcript levels as a proxy for receptor abundance at the membrane— noting that mRNA generally correlates with protein and surface expression. The selective increase in the long isoform of prolactin receptor suggests a mechanistic explanation for the elevated pSTAT5 in ACx during fatherhood. Thus, fatherhood-associated plasticity in the ACx is likely mediated by increased prolactin receptor expression rather than changes in ligand availability.

### Prolactin modulates temporal discrimination of pup USV in fathers

Does prolactin signaling contribute to the physiological plasticity observed in the ACx of fathers? To address this, we used a pharmacological approach to acutely manipulate prolactin signaling of fathers (at postnatal day 5) while performing *in vivo* two-photon calcium imaging in response to USV and their corresponding NBN.

After initial baseline imaging (**Fig. 6a** – vehicle; blue), we administered bromocriptine, a dopamine agonist that suppresses prolactin release. Then, we re-imaged the same neurons (**Fig. 6a** – green). Subsequently, we injected prolactin to determine whether it could rescue the observed effects (**Fig. 6a** – turquoise).

**Figure 6.**
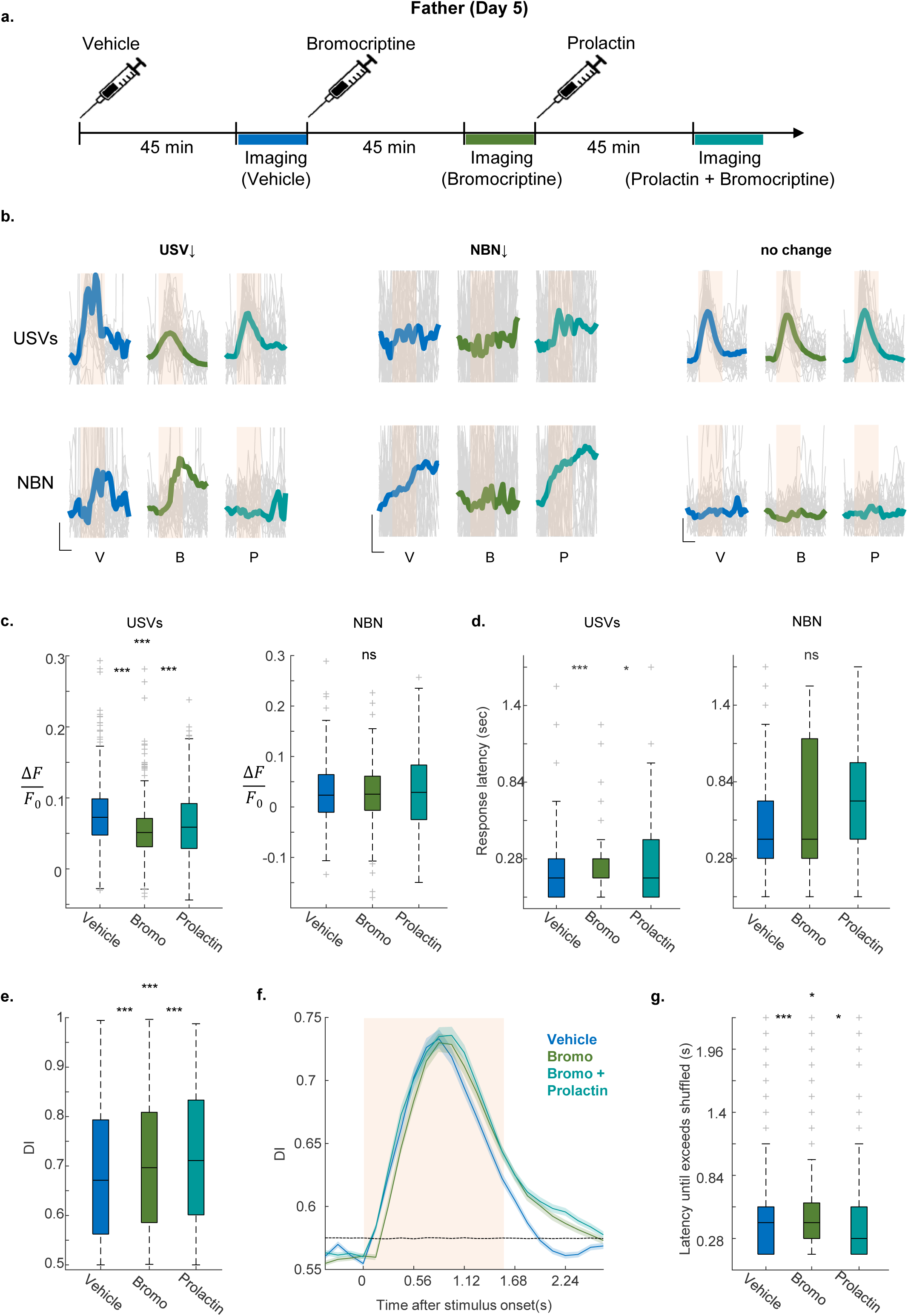
Prolactin modulates temporal discriminability of pup USVs in fathers. (a) Experimental timeline of the imaging experiment. Neuronal responses were monitored following vehicle, bromocriptine, and prolactin injections. (b) Example calcium ΔF/F traces for both USVs and NBN responses from representative neurons across treatments. Left neuron: decreased responses to USV; Center neuron: Decrease in NBN responses; Right neuron: no change. (c) Average response for both USVs and NBN. Bromocriptine induced significant suppression in response, which was partially rescued for USVs by prolactin injection. P-values USVs: V vs. B: 2.8e-53, B vs P: 1.34e-10, V vs. P: 1.1e-21. Paired LME followed by FDR for multi comparison correction (n = 376 neurons from 7 mice). P-values NBN: V vs. B: 9.2e-9, B vs P: 5.5e-7, V vs. P: 0.38. Paired LME followed by FDR for multi comparison correction (n = 237 neurons from 7 mice). (d) Latency of response to USV increased after Bromocriptine administration and restored by Prolactin, while latency of response to NBN was not significantly affected. Pval: V vs. B: 0.0001, B vs. P: 0.0119, V vs P: 0.14. Paired LME followed by FDR for multi comparison correction. For USV: n = 300, 292, 283 responsive neurons for V, B and P, respectively. or NBN: n = 56, 29, 86 responsive neurons for V, B and P, respectively. The data was obtained from 7 mice. (e) Total discriminability of USV from NBN was not impaired following prolactin levels manipulation (n = 497 neurons (responsive for USVs or NBN), from 7 mice). P values: V vs. B: 6.3351e-4, B vs P: 6.3351e-4, V vs. P: 2.31e-11. Paired LME followed by FDR for multi comparison correction. (f–g) Bromocriptine slowed discriminability of USVs from NBN; PRL restored decoding speed (n = 497 neurons (responsive for USVs or NBN, at least in one treatment), from 7 mice). P values: V vs. B: 3.55e-5, B vs P: 0.03, V vs. P: 0.03. Paired LME followed by FDR for multi comparison correction.

Administering bromocriptine induced heterogeneous changes, yet it had greater effects on responses to USV as compared to NBN **(Fig. 6b, Extended Data Fig 8a-b**). On average, bromocriptine treatment induced a modest selective reduction in neuronal responses to USV (mean reduction of 23% in response amplitude), but not to NBN (**Fig. 6c, Extended Data Fig 8a-c**). Notably, subsequent prolactin injection selectively restored neuronal responses to USVs (**Fig. 6c)**. In addition, we found a selective increase in USV response latency after bromocriptine treatment (average change at the population level: 38ms), whereas NBN latencies did not significantly change (**Fig. 6d**); this is consistent with our finding that fatherhood accelerates USV responses (**Fig. 2k**). Finally, we analyzed whether decreased prolactin levels would impair neuronal discriminability. Average single-neuron discriminability was not affected by bromocriptine proper. Paradoxically, it increased continuously during the experimental protocol (**Fig. 6e**).

Prolactin signaling also affected temporal decoding dynamics: Bromocriptine significantly slowed neuronal discrimination by 100ms, and this effect was reversed by prolactin (**Fig. 6f, g**). The change in discrimination speed cannot be explained by differences in the fraction of neurons that reached discrimination threshold (**Extended Data Fig. 8d**). Notably, this effect was specific to USV, as we found no change in temporal discriminability for WCs following either bromocriptine or prolactin treatment (**Extended Data Fig. 8e, f**).

The joint increase in single-neuron discriminability and the earlier discrimination onset (∼295ms; **Fig. 3b–d,f**) indicates that fatherhood sharpens the temporal separation of USVs from other sounds in ACx. This framing naturally directs attention to mechanisms that shift response timing, e.g., cell-intrinsic excitability changes and hormone-dependent signaling, in addition to those that simply boost amplitude.

## Discussion

To meet the demands of caregiving, prospective and new parents undergo pronounced behavioral transitions. These changes are supported by a complex array of physiological adaptations and plastic changes across multiple brain regions. While several neural circuits implicated in caregiving behavior – particularly hypothalamic circuits – have been studied in detail ^3–5,62^, parenthood is also accompanied by plasticity in primary sensory regions (e.g., olfactory and auditory ^9–20,24–27,63^).

Changes in sensory areas are thought to enhance the processing of offspring-related cues, rendering them more salient and thereby influencing parental decision-making ^24,33^. We focused on fathers in this study, which complements the existing knowledge base that is heavily biased toward maternal models. We found that––similar to motherhood––fatherhood is accompanied by plasticity in the ACx. Nevertheless, we show that the underlying effects and biological mechanisms differ between mothers and fathers.

A stereotypical and robust parental behavior is increased sensitivity to pup cries ^28,44,64,65^. In our study, cortical neurons in fathers showed improved discrimination of pup USV from their matched NBN controls. These findings contrast with previous observations in mothers, where neuronal responses in primary ACx become, in fact, less discriminative. In mothers, improved discriminability appears only downstream to primary ACx, in areas like the temporal association cortex ^15^. Additionally, in mothers, neurons in ACx generalize their responses both to spectral differences (USV vs NBN) and temporal variations (different ISI’s) of pup calls ^15,52^. In contrast, ACx neurons of males exposed to the same protocol did not generalize. If anything, they showed higher specificity to the prototype USV.

Another key difference lies in the hormonal basis of ACx plasticity. In females, oxytocin contributes to changes in sensory areas ^24,32,33,35^, while in males it does not ^37^. Instead, we identify prolactin as an important modulator of plasticity in the ACx. Reducing prolactin signaling slows discrimination while sparing accuracy, consistent with ACx plasticity in fathers primarily accelerating auditory processing rather than expanding representational capacity. This sex-specific recruitment of different neuromodulators underscores that paternal and maternal brains rely on distinct molecular routes, even in cortical pathways. Seemingly, these routes lead to functionally similar behavioral outcomes. Indeed, while common behaviors are often emphasized, the differences likely arise from divergent baseline states, expertise, and caregiving roles ^66–69^.

What underlies the improved neuronal discriminability of USV in the ACx of fathers? Since NBN-responsive neurons did not change, we interpret this result as fatherhood-sharpened ACx coding through a dual mechanism: neurons tuned to USV both increase their activity and respond with shorter latencies. This differential modulation enhances the contrast between USV and NBN. The higher and accelerated USV responses are consistent with local plastic mechanisms. This is supported by increased excitability of ACx and the elevated prolactin signaling, which both point to cell-intrinsic plasticity ^43^. The delayed response latency to USV following manipulation of prolactin levels in fathers further supports this view. We do not exclude additional circuit mechanisms; for example, selective engagement of local inhibitory subtypes that sculpt temporally patterned inputs could sharpen USV responses while leaving NBN averages unchanged^70–74^. Beyond local plasticity, top-down regulation from higher-order regions may also be involved. For instance, mechanisms like modulation by the paraventricular nucleus or orbitofrontal cortex observed in females, may also play a role in males ^33,75^.

We show that fatherhood-related neuronal changes are correlated with elevated prolactin signaling in the ACx (**Fig. 5**). Unlike biparental species such as California mice, the increased prolactin signaling here does not result from elevated circulating prolactin levels ^76^. Instead, we found that an elevation in prolactin receptor level mediates this increase (**Fig. 4d,e**) and likely accounts for its specificity. We cannot exclude that other modulators of prolactin signaling, like the levels of intracellular regulators (e.g., SOCS1, SOCS3) or STAT5 itself, which can also contribute to the elevated signal detected in the ACx of fathers ^77,78^.

Is the elevated excitability of pyramidal neurons during fatherhood related to increased prolactin signaling? We show that during fatherhood, the ACx neurons have increased input resistance, likely underlying the increase in neuronal excitability^79^. Moreover, we found that these neurons also show a depolarized resting membrane potential, which can account for an increase in thresholds and a decrease in the rate of rise and amplitude of the action potential. Notably, both increase in input resistance and membrane depolarization might result, for example, from a decrease in the potassium conductance ^80^. In line with our results, prior work shows that prolactin receptor activation can rapidly re-tune intrinsic neuronal properties via inhibition of BK Ca²⁺-activated K⁺ channels ^81^ or reduction in the transient A-type K⁺ current ^82^. It is therefore plausible, that elevated prolactin during fatherhood may reduce specific K⁺ currents in ACx pyramidal neurons, thus increasing their excitability. Targeted manipulations of prolactin signaling and K⁺ conductances *in vivo* can help test this hypothesis in future experiments.

In summary, we find that fatherhood does not simply recruit the canonical subcortical “parental” circuits; it also remodels sensory coding in its own unique way. This work broadens our understanding of plasticity in the parental brain beyond maternal models, and it highlights the role of hormones such as prolactin in sculpting sensory circuits in socially relevant contexts.

## Acknowledgements

We thank members of the Mizrahi laboratory for comments on the manuscript. We thank Gerard Elberg and Maya Groysman for technical help. This work was supported by a grant from the Israel Science Foundation to A.M (#511/23) and to A.B. (#1202/23). The work was supported by the Gatsby Charitable Foundation. B.H was supported by the Shimon Peres postdoctoral fellowship and the Edmond and Lily Safra Center for Brain Sciences. A.M. is the Eric Roland Chair in Brain Sciences, and A.B. is the Cecile and Seymour Alpert Chair in Pain Research. Figure 1B,H and Figure 2B were created using <u>biorender.com</u>.

## Author Contributions

Conceptualization: B.H., A.B., and A.M. Methodology: B.H., and A.M. Investigation: B.H., T.P., P.R.R., and H.S. Formal analysis: B.H., T.P., and P.R.R. Visualization: B.H., T.P., and P.R.R. Software: B.H., and T.P. Supervision: A.M. Funding acquisition: A.M. Writing: B.H. P.R.R., A.B., and A.M.

## Competing Interests

The authors declare that they have no competing interests.

## Data availability

The source data that support the findings of this study will be deposited in the Zenodo repository (Resource Type: Dataset) and will be available under the DOI: 10.5281/zenodo.17396987. These data will be publicly released upon publication. Additional raw data are available from the corresponding author upon reasonable request.

## Extended Data Figure Legends

**Extended Data Figure 1.** Ethograms of post-mating males and fathers. Representative ethograms for individual mice at Post-mating (a) and Father (b) time points (n = 9 mice). Colors denote behavioral categories: self-grooming (gray), pup sniffing (yellow), nest building (brown), pup grooming (blue), crouching (purple), and pup retrieval (green). Each trace spans the entire session duration.

**Extended Data Figure 2.** Isolated males after fatherhood exhibit reduced parental care. (a) Experimental timeline depicting the three assessment time points: Post-mating (3–5 days after mating), Father (4–6 days after parturition) and Post fatherhood (7 days after separation from dam and pups). (b-c) Fraction of time devoted to self-grooming, sniffing and nest building (b) and pup grooming and crouching (c) for post mated males and fathers (n = 18,18 and 11 for post mated males, fathers and post-fatherhood males, respectively). (d) Proportion of mice exhibiting pup retrieval behavior for across the three time points (n = 18 mice). (e) Total time dedicated for pup retrieval, latency to retrieve first pup and pup retrieval span (time from retrieval of first to the last pup) for Post-mated, Father and Post fatherhood males (n = 18,18 and 11, respectively). (f) Proportion of mice exhibiting pup retrieval behavior for Post-mated, Father and Post fatherhood males assessed longitudinally (n = 5 Mice, same mice for all time points). Data are median ± IQR; comparisons by Kruskal–Wallis test with Tukey-Kramer’s post hoc correction; ns *p* > 0.05, \**p* < 0.05, \*\**p* < 0.01, \*\*\**p* < 0.005.

**Extended Data Figure 3.** Fatherhood alters the salience of pup vocalizations. (a) Difference in sound intensity (RMS amplitude) between USV and NBN speakers across the arena. (b) Individual mouse heatmaps illustrating spatial preference change (sound – habituation) for mice as post mated males and as fathers (n = 13 mice). (c) Dynamics of spatial preference for mice as post mated males and as fathers (n = 13 mice). (d) Total distance mice traveled during 5 minutes of habituation and during sound exposure for mice as post mated males and as fathers (n = 13 mice). (e) Number of arena midline crossings during habituation and sound exposure (n = 13 mice). Data are median ± IQR; comparisons by Kruskal–Wallis test with Tukey-Kramer’s post hoc test and Wilcoxon signed-rank test; ns *p* > 0.05, \**p* < 0.05, \*\**p* < 0.01, \*\*\**p* < 0.005.

**Extended Data Figure 4.** Best-frequency distribution shifts during fatherhood. (a) BF distribution of 3 mice. pvals: 1.06e-6, 9.1584e-7 and 1.125e-17, for mouse #1, –3, respectively (Wilcoxon signed-rank test). (b) Histogram (gray bars) shows the probability density of mean ∣ΔBF∣ values obtained from 2,000 bootstrap resamples (with replacement) of the pooled absolute tuning-shift magnitudes (n = 295 neurons, 3 mice). The red line marks the observed mean ∣ΔBF∣ in the actual data. Dashed black lines mark the 2.5th and 97.5th percentiles of the bootstrap distribution, defining a 95% confidence interval. Bootstrap analysis confirms that the average magnitude of tuning-curve shifts is reliably greater than zero. (c). Fraction of neurons responsive to USVs and NBN (upper and lower rows, respectively) across the three time points. Each point represents a mouse. (d). Fraction of neurons responsive to all protocol stimuli across the three time points. Each point represents a mouse. Data were obtained from 9 mice. Values are median ± IQR; *p<0.05, **p<0.01, **p<0.005 by LME followed by FDR for multi comparison correction.

**Extended Data Figure 5.** Significant correlation between total discriminability and speed. No improvement in discrimination of wriggling calls (WCs) in fathers. (a) Transient decrease in response latency to USVs following fatherhood, but not to NBN (n= neurons, from 9 mice). P-values: PM vs. F: 6.1743e-6. PM vs. PF: 0.14, F vs. PF: 0.0063. LME followed by FDR for multi comparison correction. (b) Strong correlation between increased discriminability and decreased latency. P-values: 4.8e-6, 1.9e-10, 0.0029 for post mating, father and post-fatherhood, respectively. (c-d) Time-resolved decoding analyses revealed faster discrimination of WCs during fatherhood (n= 388, 304 and 169 neurons (responsive to WCs or their corresponding NBN), from 9, 9 and 6 mice for the different time points).

**Extended Data Figure 6.** Action-potential properties. (a-d). Pyramidal neurons in fathers exhibit an increase in AP threshold potential (a), a decrease in AP amplitude (b) and maximal rate of rise of AP (dV/dt_max_) (c), without changes in AP half-width (d). P-values: 9.1835e-04, 0.029, 0.015 and 0.76, for comparisons in A-D, respectively (Unpaired t-test).

**Extended Data Figure 7.** Prolactin signaling is enhanced in auditory but not visual cortex or PAG during fatherhood. (a). Quantification of pSTAT5+ cell density across multiple brain regions shows increased prolactin signaling in ACx and medial preoptic area (mPOA), but not in periaqueductal gray (PAG) or visual cortex. MPOA: One-way ANOVA: 0.0005. PM vs F: 3e-4, F vs PF: 0.06, PM vs PF 0.17, after Tukey-Kramer multiple comparison correction. Data were collected from 5 brain slices per mouse, across 5 mice. PAG: One-way ANOVA: 0.17. Data were collected from 5 brain slices per mouse, across 4 mice. VCx: One-way ANOVA: 0.94. Data were collected from 5 brain slices per mouse, across 4 mice. Data are median ± IQR; *p<0.05, **p<0.01, **p<0.005 by One-way ANOVA followed by Tukey-Kramer correction for multiple comparisons.

**Extended Data Figure 8.** Additional data from pharmacological manipulation. (a–b) Proportion of significantly affected neurons following prolactin levels manipulation, shown for all neurons and for responsive neurons only. (c) Fraction of responsive neurons remained stable across treatments. (d) Decoding improvements were not explained by differences in the proportion of neurons below threshold after each treatment. (e–f) Temporal discrimination effects were absent for WCs. P values f: V vs. B: 6.3351e-4, B vs P: 6.3351e-4, V vs. P: 2.31e-11. LME followed by FDR for multi comparison correction. Data were obtained from 8 mice. Values are median ± IQR; *p<0.05, **p<0.01, **p<0.005 by LME followed by FDR for multi comparison correction.

## Methods

### Animals and surgical procedures

We used ICR:CD1 mice for the study (Envigo Israel). All experimental procedures were conducted in accordance with the Hebrew University Animal Care and Use Committee (NS-16966, NS-17642, NS-17827). We used a virus expressing a calcium indicator in the cytoplasm and a red fluorescent protein in the nucleus (AAV9-hsyn-GCaMP6s-P2A-nls-dTomato) that was produced at The Edmond and Lily Safra Center for Brain Sciences virus core facility (https://elsc.huji.ac.il/research-and-facilities/expertise-centers/elsc-vector-core-facility/) and was assessed at a titer of 10^12^. The virus (200 nl) was injected using NanoJect 2 to the ACx of the left hemisphere of juvenile mice (age range, postnatal day 26 (P26) to P30). The injection site was sealed with bonewax. During the same procedure, a head bar was fixed to the top of the skull using dental cement. A 3 mm chronic glass window was implanted above the injection site according to published protocols ^83^ at 14–28 d after virus injection. Both procedures were performed under 2% isoflurane anesthesia. The hair was initially removed from the surgery area using a commercial hair removal cream and rinsed with rubbing alcohol. Lidocaine was injected under the skin as a local analgesic. Mice were injected subcutaneously with carprofen (4 mg/kg) during each procedure.

### Timeline

Throughout the study, the experiments were conducted on mice at three time points: (1) 3-5 days after mating, (2) 4-6 days after parturition, and (3) 7-8 days post-isolation (i.e., post-mating males, fathers, and post-fatherhood males, respectively). Males were cohabited with the female and their pups until isolation. For the third experimental group, the males were isolated once the pups matured – between P14-P21.

### Paternal behavior assessment

Male mice were recorded alone in their home cage for 30 minutes – 15 minutes of habituation followed by 15-minute, during which pups were placed on the opposite side of the cage to elicit pup retrieval. The videos were analyzed and manually annotated using EthoVision XT16, for parental (pup grooming, crouching and pup retrieval) and non-parental (self-grooming, pup sniffing and nest building) behaviors. Pup retrieval behavior was assessed by locating 4 pups at the opposite side of the cage. Both self and foreign pups were retrieved. We calculated the total time, latency to begin (1^st^ pup) and time span (from 1^st^ pup to 4^th^ pup) of the behavior.

### Auditory stimuli preparation

ICR:CD1 pup USV were recorded with a one-quarter inch microphone (Brüel & Kjær) from P4–P5 pups (250 kHz sampling rate). Additional details can be found in our previous paper ^14^. USV consisted of 7 syllables. To synthesize NBN, we produced a narrow band noise at corresponding frequency range, along the time window for each syllable in the USV. Within each syllable window, the NBN segment was scaled to match the root-mean-square (RMS) amplitude of the corresponding recorded USV syllable, and after adding background noise, the entire NBN call was globally RMS-matched to the original call. In addition, 5 inter syllable interval (ISI) – modulated calls were prepared based on the original recorded prototype USV call, by introducing 75, 125, 275, 375 and 575 ms spacing between the syllables (prototype USV call ISI was ∼175 ms). Background noise was added between the syllables of all synthetic stimuli to mimic the prototype call.

### Auditory protocol

The auditory protocol consisted of USV, their corresponding NBN, WCs, their corresponding NBN and the ISI-modulated USVs, as mentioned before. The speaker was driven at a 250 kHz sampling rate via a driver (model ED1, TDT). The stimuli order was pseudorandomized. Inter-stimulus intervals (ISIs) were drawn on each trial from a uniform distribution centered at 3.5 s with ±0.2 s jitter.

To detect neuronal tuning, we generated 14 log-spaced pure tones with frequencies ranging from 4 to 64 kHz. The sequence of the tones was randomized. The speaker was driven at a 500 kHz sampling rate via a driver (model ED1, TDT). Sound intensity was calibrated to 75 ± 2 dB SPL for all presented sound frequencies.

### Auditory-driven spatial preference

Male mice were placed in a square arena equipped with a free-field speaker (model ES1, TDT) on each of two opposing walls. The mice were recorded for 30 minutes – 15 minutes of habituation followed by 15-minute, during which recorded ICR pup ultrasonic vocalizations and their corresponding NBN was played through the speakers. Each sound was continuously repeated with inter-bout intervals drawn uniformly from ISI ± 1 s (default ISI = 5 s). The speaker was driven at a 500 kHz sampling rate via a driver (model ED1, TDT). Sound intensity was calibrated to 75 ± 2 dB SPL (a fixed digital attenuation of 10 dB was applied equally to both channels) for all presented sound frequencies.

Using EthoVision XT16, the arena was divided into 3 parts (2 peripheral and one neutral central zone). Mouse position was tracked as time-stamped (x,y) coordinates with binary zone labels for the two speaker zones (USV, NBN). Brief tracking dropouts or spikes were flagged by a step-distance threshold and NaNs, then linearly interpolated. Zone traces were median-filtered and linearly gap-filled. Spatial occupancy heatmaps were computed by kernel density estimation on a 100×100 grid for visualization. Preference was quantified per mouse as a zero-centered Preference Index (PI), comparing the NBN/USV time ratio during the sound period versus habituation:

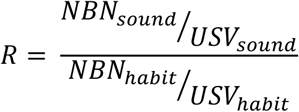

Then set:

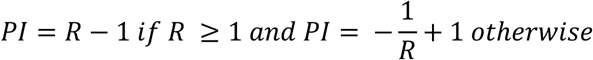

Meaning PI = 0: no change; PI >,0: shift toward NBN; PI < 0: shift toward USV. Time-resolved PI was also calculated using cumulative windows (0.5-min steps) and fixed 1-min bins. Locomotion controls included total path length and midline crossings.

The sound intensity and distribution throughout the arena was validated using a one-quarter inch microphone (Brüel & Kjær).

### Two-photon calcium imaging

We imaged GCaMP6s-labeled neurons in layer 2/3 using a custom-built^84^ galvo-mirror scanning two-photon microscope with a frame rate of 7.2 Hz. Two-photon excitation (950 nm) was delivered through a DeepSee femtosecond laser (Mai Tai, SpectraPhysics). Imaging was performed through a water-immersion objective (0.8 numerical aperture; model CF175, Nikon) and detected through GaAsP Photomultiplier Tubes (Hamamatsu). The imaging field size was set to 260×260 microns over a 512×210 pixel window (the pixel size was chosen to maximize imaging rate using our galvo-galvo scan head). We used Scanimage software ^85^ for acquisition and online drift correction (using the red channel).

### Calcium imaging analysis

Image stacks were corrected online for motion using the red channel as reference. Regions of interest (ROIs) were selected manually for each cell, for all blocks taken together. We ensured that all selected neurons were within the field of view throughout all blocks. Raw fluorescence time series (F(t)) were obtained for each cell by averaging across pixels within each ROI. Baseline fluorescence (F_0_) was computed by taking the mean F(t) prior to each stimulus (1 sec). The change in fluorescence relative to baseline, ΔF/F_0_, was computed as (F(t)-F_0_)/F_0_. A neuronal response to a single trial was calculated as the average of ΔF/F_0_ of that neuron, 1 s following sound onset. A neuron was classified as responsive if its responses (average of 1 s after stimulus for each trial) were significantly higher than the pre-stimulus signal (Wilcoxon signed rank test, p < 0.05). All analyses were performed in custom-written MATLAB code and will be available for download.

*Response latency*: A threshold was set for each neuron as the pre-stimulus response plus three standard deviations of the pre-stimulus response. Latency was the index of the first post-stimulus time point at which the response exceeded this threshold. Only neurons classified as responsive for that stimulus (see above) were included.

*Lifetime sparseness*: we computed lifetime sparseness across the 14 tone frequencies in the tuning set for each neuron:

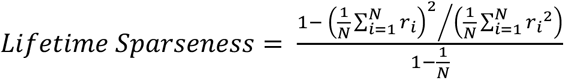

Where N is one of the 14 tone frequencies in the tuning set, and r is the mean response of the neuron to each stimulus. Values were rectified to zero. Analyses were restricted to responsive neurons, defined as cells that met our responsiveness criterion for at least one of the 14 frequencies.

### Best frequency analysis

BF was defined as the tone frequency (from the 14-frequency set) eliciting the largest trial-averaged ΔF/F during the first second after stimulus onset. Only responsive neurons to at least one frequency were analyzed.

Per-mouse directionality of BF shifts:

Within each mouse, we computed signed BF shifts as the difference of BF at fatherhood and at post mating stage (in frequency-index units, 1–14). To test whether shifts were directionally biased, we (i) calculated the observed mean change in BF per mouse; (ii) generated a sign-flip permutation null by randomly multiplying each neuron’s change in BF by ±1 and recomputing the mean over 10,000 permutations; and (iii) obtained a two-tailed permutation *p*-value as the fraction of null means with absolute value ≥ ∣ΔBF∣. For visualization, kernel density estimates (KDEs) of the null and actual distributions were plotted on a shared grid, with vertical markers at zero and average change in BF.

To assess whether the magnitude of retuning exceeded chance, we pooled neurons across mice and computed ∣ΔBF∣. A shuffle-pairings null was constructed by breaking the within-neuron correspondence between stages: BF values from fatherhood stage were randomly permuted across neurons and paired with each neuron’s post mating stage BF. This shuffle was repeated 10,000 times to form a null distribution. We compared the observed and null distributions by a permutation test on the mean of |ΔBF| under the shuffled (null) distribution, using the 10,000 null means (reporting the tail probability that the null mean ≥ the observed mean). A bootstrap (2,000 resamples) provided a 95% confidence interval for the observed mean ∣ΔBF∣. KDEs of the observed and null distributions were plotted for display.

*Discrimination Index*: To measure how well neurons discriminate between stimuli, we calculated a receiver operating characteristics (ROC) curve between the distributions of each neuron’s responses to these sounds and calculated its area under the curve (AUC). When the numbers of USV and NBN trials differed, we balanced sample sizes by random down-sampling to the smaller set and averaged DI across 100 resamplings. An AUC value close to 0.5 indicates low discrimination, whereas an AUC value away from 0.5 indicates high discrimination. To compensate for the neuronal preferences of either sound, we calculated the discrimination index (DI):

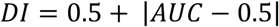

We calculated two forms of DI values: between the USV (or WCs) and the corresponding NBN stimuli at each stage, and between different stages for a given stimulus.

*Discrimination Index* dynamics: we computed DI in a sliding window of 0.42 sec across the peri-stimulus epoch to obtain a DI time course per neuron and stage. When the numbers of USV and NBN trials differed, we balanced sample sizes by random down-sampling to the smaller set and averaged DI across 100 re-samplings. Only neurons that were responsive to USV or NBN were included. To define a data-driven “discriminability present” boundary at each time bin and stage, we generated a shuffle-null by randomly permuting USV/NBN labels within neuron and recomputing DI, and set the per-bin threshold to the shuffle mean + 3 SD. Then, the “DI latency” was defined as the earliest post-stimulus time bin at which the DI time course exceeded the stage-specific shuffle threshold for ≥ 3 consecutive frames (to ensure temporal stability). Neurons that never crossed threshold were assigned NaN and were excluded from latency distributions but were summarized per mouse as the fraction of neurons “below threshold”.

### pSTAT5 immunohistochemistry

Anesthetized mice were given an overdose of Pental and were perfused transcardially with phosphate-buffered saline (PBS) followed by 4% paraformaldehyde (PFA) in PBS. Brains were postfixed for 12–24 h in 4% PFA in PBS and then cryoprotected for >24 h in 30% sucrose in PBS. Coronal slices of 50 µm were made using a freezing microtome (Leica SM 2000R) and preserved in PBS.

Prior to immunofluorescence staining, antigen retrieval was performed by incubating sections for 15 minutes in citric acid (pH 6) at 80°C, followed by 10 minutes incubation in Hydrogen Peroxide 3%. Sections were then washed with PBS and incubated in blocking solution (1% Goat Serum, 0.1% Triton in PBS) for two hours. After PBS wash, sections were incubated in rabbit pSTAT5 primary antibody (pSTAT5 Tyr 694, Cat#: C11C5, 1:500; Cell Signaling Technology) for 72 hours at 4°C, followed by two-hours incubation in goat anti-rabbit Cy5 conjugated secondary antibody (1:250; Jackson ImmunoResearch, Cat #111-175-144, RRID: AB_2338013). Then, the slices were washed with PBS and incubated in 2.5 µg/ml of DAPI (Santa Cruz, Cat #sc-3598) in PBS for 10 minutes. Finally, the slices were washed with PBS and mounted onto slides and cover slipped with mounting media (Vectashield H-1200).

Sections were imaged using an Olympus IX-81 epifluorescent microscope with a 10× objective lens (0.3 NA; Olympus). Brain areas were determined according to their anatomy using Allen Brain Atlas and counting was done manually by marking a region within the brain area of interest and then dividing the counted signal points by the region area.

### Auditory cortex prolactin level quantification

Auditory cortex was identified, cut out with a 2.5 mm punch and immediately freeze in –80°C. We used Mouse Prolactin ELISA kit (#EMPRL, *Invitrogen*), for quantifying prolactin level. Before use, the samples were thawed at 4°C for 12 hours, and then manually homogenized using 30 G needle in 100 µl of Diluent C until the sample appears well-homogenized. Samples’ Prolactin concentration was normalized to samples’ weight.

### RT-PCR

Mice were given an overdose of Pental and brains were extracted. Auditory cortex was identified, cut out with a 2.5 mm punch and immediately freeze in –80°C. RNA was produced using NucleoSpin RNA XS, Micro kit for RNA purification (MACHEREY-NAGEL, Cat#: 740902.50), following the manufacturer’s protocol. RNA quality and concentration were determined by measuring the A260/280 and A260/230 absorbance ratios using the NanoDrop spectrophotometer (Thermo Fisher Scientific). Subsequently, 500 ng of RNA was reverse transcribed into cDNA using Maxima H Minus Reverse Transcriptase (200 U/μL, Thermo Fisher Scientific, Cat# EP0751). qPCR procedures were carried out at the Center for Genomic Technologies, Hebrew University of Jerusalem. Each sample was quantified in triplicates for the following targets and housekeeping gene (GAPDH) as an internal control.

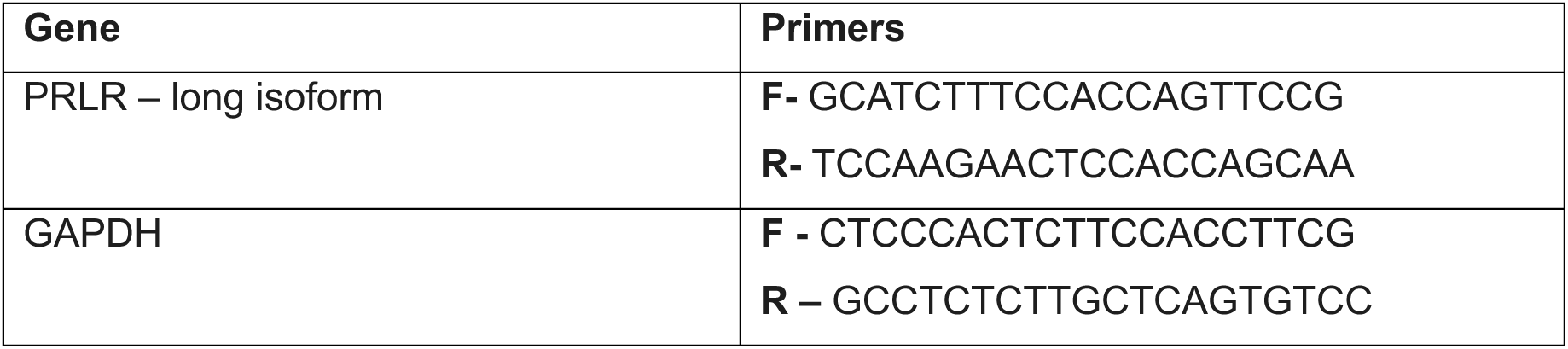

### Pharmacological intervention

The following protocol was used on male mice 5 days post-partum (overall similar to Smiley et al 2022). First, mice were injected with vehicle solution i.p. (70% saline/ 30% ethanol), followed by an imaging session 45 minutes after injection. Then, mice were injected i.p. with Bromocriptine (200 µl, 7.78 mg/kg, 30% ethanol), followed by an imaging session 45 minutes after injection. Finally, mice were injected with Prolactin i.p. (200 µl, 0.05 mg/kg), followed by an imaging session 45 minutes after injection.

### Electrophysiology

#### Acute brain slice preparation

For all electrophysiological recordings, we used age-matched male mice categorized into two groups: post-mating males and fathers. On the day of the experiment, mice were taken out from their home cage and anesthetized using ketamine/domitor mixture (dosage). Deep anesthesia was validated by a loss of toe-pinch reflex. Once fully anesthetized, mice were perfused transcardially with 20 mL oxygenated ice-cold sucrose-based cutting solution containing (in mM): 110 sucrose, 5 D-glucose, 120 NaCl, 56 NaHCO_3_, 6 KCl, 2.5 NaH_2_PO_4_, 14 MgCl_2_, and 1 CaCl_2_. Brains were quickly removed and immersed in the cutting solution. Coronal brain sections (300 μm thick) containing primary auditory cortex (ACx) region were prepared using a Leica VT1200S vibrating blade microtome (Leica Biosystems) and collected in ice-cold cutting solution. The slices were then transferred to warm artificial CSF (aCSF) containing (in mM): 25 D-glucose, 250 NaCl, 50 NaHCO_3_, 5 KCl, 2.5 NaH_2_PO_4_, 2 MgCl_2_, and 4 CaCl_2_, maintained at 34°C for 30 min to allow recovery. Following initial recovery period, slices were left in aCSF at room temperature (RT, ∼24°C) for at least 60 min before transferring to the recording chamber for electrophysiological recordings. Throughout the procedure, all solutions were continuously bubbled with carbogen (95% O_2_/5% CO_2_).

#### Whole-cell patch clamp

Brain sections containing layer 2/3 pyramidal neurons of the primary auditory cortex (ACx) were visualized with infrared differential interference contrast (IR-DIC) microscopy. These neurons were identified by their large pyramidal shape and regular firing pattern. All electrophysiological recordings were carried out at RT.

The recording electrodes were pulled from borosilicate glass capillaries (1403306; Hilgenberg, Germany) using an electrode puller (P-1000; Sutter Instruments, Navato, CA) and had a resistance of 3-5 MΩ. These were back-filled with K-gluconate-based intracellular solution containing (in mM): 135 K-gluconate, 6 NaCl, 2 MgCl_2_, 10 HEPES, 0.2 EGTA, 2 MgATP, and 0.3 Na_3_GTP, with an osmolarity of 280 mOsm and pH adjusted to 7.25 with KOH. The seal was ruptured after the cells reached resistance of >2 GΩ. Once in whole-cell mode, we waited for at least 5 min for seal stabilization, and diffusion of the internal solution before acquiring intrinsic excitability properties in current clamp mode. The pipette series resistance (R_s_) was continuously monitored and recordings were excluded from analysis if R_s_ of the neurons was >20 MΩ and/or resting membrane potential (RMP) > −60 mV. Recordings were sampled at 50 kHz, digitized using a Digidata 1550B apparatus (Molecular Devices, San Jose, CA), filtered at 10 kHz, and amplified using a Multiclamp 700B amplifier. The data were analyzed off-line by Clampfit 11.2 (Molecular Devices, San Jose, CA).

#### Recording and analysis of neuronal excitability properties

All recordings were carried out at the natural resting membrane potential (RMP) of the neurons. No currents were applied to modify the membrane potential.

*Resting membrane potential* was recorded within 3-5 min after breaking the seal.

The *neuronal firing* was studied by applying a series of depolarizing 1s pulses starting from 50 pA to a maximum of 300 pA in 50 pA increments. Then the *f*-I relationship between applied stimuli and the frequency of AP firing was established.

To assess neuronal gain, a linear function was fitted to the *f*-I curve, and the slope (*m*) of the fit was taken as the gain.

*Input resistance* (*R_in_*) was determined from the steady-state voltage response of the neuron to a hyperpolarizing current injection of –50 pA and linearly fitting the voltage change to the injected current.

For studying the rheobase, AP threshold, AP amplitude, half-width, and rates of changes of membrane voltage, single APs evoked by injecting square current pulses for 10 ms in 10 pA steps, were analyzed.

*Rheobase* (threshold current) was defined as the minimum amount of current needed to induce the first AP.

The *maximum rate of rise (dV/dt_max_)* was determined from the maximum dV/dt value on the rising phase of the first derivative trace of AP.

*AP threshold* was defined as the membrane potential at which the first derivative of the voltage trace (*dV/dt*) reached 30 V/s.

*AP amplitude* was measured from the threshold to the peak of the AP.

*AP half-width* was calculated as the AP duration at the half-maximal amplitude. All analyses were performed by the experimenter blind to the group.

### Statistical analysis

All statistical analyses were performed in MATLAB. We used the sign-rank test, the Wilcoxon rank-sum test, and the student’s t-test, as indicated in the text. To control for random effects, we used Linear mixed-effects modeling as described below. In cases of multiple comparisons, *P* values were controlled using Benjamini–Hochberg FDR.

#### Linear mixed-effects modeling

For Group comparisons for neuron-level outcomes were tested with linear mixed-effects (LME) models. For unpaired analyses we fit

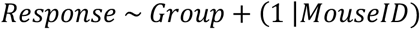

treating Group (e.g., Stage) as a fixed effect and MouseID as a random intercept to account for within-mouse dependence. When the same neurons were tracked across conditions, we used a paired model with an additional random intercept for Cell-ID (CellIDs globally unique across the dataset):

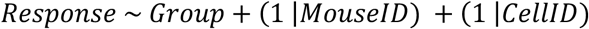

Models were estimated by Restricted (Residual) Maximum Likelihood. For outcomes with non-Gaussian distributions, we applied a rank-based approach by replacing the response with its tied ranks prior to fitting.

Planned pairwise contrasts between Group levels were tested as Wald tests on the fixed-effect coefficients. Where multiple contrasts were evaluated, *P* values were controlled using Benjamini–Hochberg FDR.

## Data availability

The data supporting the findings of this study are available from the corresponding author upon reasonable request.

## Code availability

Custom MATLAB code used for behavioral, imaging, and electrophysiological analyses will be made available upon publication or upon reasonable request from the corresponding author.

## Notes

### Competing Interest Statement

The authors have declared no competing interest.

## References

1. Kölliker, M. (2012). The Evolution of Parental Care N. J. Royle and P. T. Smiseth, eds. (Oxford University Press) 10.1093/acprof:oso/9780199692576.001.0001.

2. van Noordwijk, M.A., and van Schaik, C.P. (2005). Development of ecological competence in Sumatran orangutans. Am J Phys Anthropol 127, 79–94. 10.1002/ajpa.10426.

3. Inada, K., Hagihara, M., Tsujimoto, K., Abe, T., Konno, A., Hirai, H., Kiyonari, H., and Miyamichi, K. (2022). Plasticity of neural connections underlying oxytocin-mediated parental behaviors of male mice. Neuron 110, 2009–2023.e5. 10.1016/j.neuron.2022.03.033.

4. Majdoubi, M.E.I., Poulain, D.A., and Theodosis, D.T. (1996). The glutamatergic innervation of oxytocin– and vasopressin-secreting neurons in the rat supraoptic nucleus and its contribution to lactation-induced synaptic plasticity. European Journal of Neuroscience 8. 10.1111/j.1460-9568.1996.tb01600.x.

5. Wu, Z., Autry, A.E., Bergan, J.F., Watabe-Uchida, M., and Dulac, C.G. (2014). Galanin neurons in the medial preoptic area govern parental behaviour. Nature 509. 10.1038/nature13307.

6. Champagne, F., Diorio, J., Sharma, S., and Meaney, M.J. (2001). Naturally occurring variations in maternal behavior in the rat are associated with diberences in estrogen-inducible central oxytocin receptors. Proceedings of the National Academy of Sciences 98, 12736–12741. 10.1073/pnas.221224598.

7. Ribeiro, A.C., Musatov, S., Shteyler, A., Simanduyev, S., Arrieta-Cruz, I., Ogawa, S., and Pfab, D.W. (2012). siRNA silencing of estrogen receptor-α expression specifically in medial preoptic area neurons abolishes maternal care in female mice. Proceedings of the National Academy of Sciences 109, 16324–16329. 10.1073/pnas.1214094109.

8. Fang, Y.Y., Yamaguchi, T., Song, S.C., Tritsch, N.X., and Lin, D. (2018). A Hypothalamic Midbrain Pathway Essential for Driving Maternal Behaviors. Neuron 98. 10.1016/j.neuron.2018.02.019.

9. Lau, B.Y.B., Krishnan, K., Huang, Z.J., and Shea, S.D. (2020). Maternal Experience-Dependent Cortical Plasticity in Mice Is Circuit– and Stimulus-Specific and Requires MECP2. The Journal of Neuroscience 40, 1514– 1526. 10.1523/JNEUROSCI.1964-19.2019.

10. Liu, R.C., and Schreiner, C.E. (2007). Auditory Cortical Detection and Discrimination Correlates with Communicative Significance. PLoS Biol 5, e173. 10.1371/journal.pbio.0050173.

11. Cohen, L., and Mizrahi, A. (2015). Plasticity during motherhood: Changes in excitatory and inhibitory layer 2/3 neurons in auditory cortex. Journal of Neuroscience 35. 10.1523/JNEUROSCI.1786-14.2015.

12. Cohen, L., Rothschild, G., and Mizrahi, A. (2011). Multisensory integration of natural odors and sounds in the auditory cortex. Neuron 72. 10.1016/j.neuron.2011.08.019.

13. Rothschild, G., Cohen, L., Mizrahi, A., and Nelken, I. (2013). Elevated correlations in neuronal ensembles of mouse auditory cortex following parturition. Journal of Neuroscience 33. 10.1523/JNEUROSCI.4656-12.2013.

14. Tasaka, G.I., Guenthner, C.J., Shalev, A., Gilday, O., Luo, L., and Mizrahi, A. (2018). Genetic tagging of active neurons in auditory cortex reveals maternal plasticity of coding ultrasonic vocalizations. Nat Commun 9. 10.1038/s41467-018-03183-2.

15. Tasaka, G.I., Feigin, L., Maor, I., Groysman, M., DeNardo, L.A., Schiavo, J.K., Froemke, R.C., Luo, L., and Mizrahi, A. (2020). The Temporal Association Cortex Plays a Key Role in Auditory-Driven Maternal Plasticity. Neuron 107. 10.1016/j.neuron.2020.05.004.

16. Haimson, B., and Mizrahi, A. (2023). Plasticity in auditory cortex during parenthood. Hear Res 431, 108738. 10.1016/j.heares.2023.108738.

17. Elyada, Y.M., and Mizrahi, A. (2015). Becoming a mother—circuit plasticity underlying maternal behavior. Curr Opin Neurobiol 35, 49–56. 10.1016/j.conb.2015.06.007.

18. Kim, P., Strathearn, L., and Swain, J.E. (2016). The maternal brain and its plasticity in humans. Horm Behav 77, 113–123. 10.1016/j.yhbeh.2015.08.001.

19. Miranda, J.A., and Liu, R.C. (2009). Dissecting natural sensory plasticity: Hormones and experience in a maternal context. Hear Res 252, 21–28. 10.1016/j.heares.2009.04.014.

20. Miranda, J.A., Shepard, K.N., McClintock, S.K., and Liu, R.C. (2014). Adult plasticity in the subcortical auditory pathway of the maternal mouse. PLoS One 9. 10.1371/journal.pone.0101630.

21. Dunlap, A.G., Besosa, C., Pascual, L.M., Chong, K.K., Walum, H., Kacsoh, D.B., Tankeu, B.B., Lu, K., and Liu, R.C. (2020). Becoming a better parent: Mice learn sounds that improve a stereotyped maternal behavior. Horm Behav 124. 10.1016/j.yhbeh.2020.104779.

22. Galindo-Leon, E.E., Lin, F.G., and Liu, R.C. (2009). Inhibitory Plasticity in a Lateral Band Improves Cortical Detection of Natural Vocalizations. Neuron 62. 10.1016/j.neuron.2009.05.001.

23. Lin, F.G., Galindo-Leon, E.E., Ivanova, T.N., Mappus, R.C., and Liu, R.C. (2013). A role for maternal physiological state in preserving auditory cortical plasticity for salient infant calls. Neuroscience 247, 102–116. 10.1016/j.neuroscience.2013.05.020.

24. Marlin, B.J., Mitre, M., D’Amour, J.A., Chao, M. V., and Froemke, R.C. (2015). Oxytocin enables maternal behaviour by balancing cortical inhibition. Nature 520. 10.1038/nature14402.

25. Royer, J., Huetz, C., Occelli, F., Cancela, J.M., and Edeline, J.M. (2021). Enhanced Discriminative Abilities of Auditory Cortex Neurons for Pup Calls Despite Reduced Evoked Responses in C57BL/6 Mother Mice. Neuroscience 453. 10.1016/j.neuroscience.2020.11.031.

26. Moreno, A., Gumaste, A., Adams, G.K., Chong, K.K., Nguyen, M., Shepard, K.N., and Liu, R.C. (2018). Familiarity with social sounds alters c-Fos expression in auditory cortex and interacts with estradiol in locus coeruleus. Hear Res 366. 10.1016/j.heares.2018.06.020.

27. Chong, K.K., Anandakumar, D.B., Dunlap, A.G., Kacsoh, D.B., and Liu, R.C. (2020). Experience-dependent coding of time-dependent frequency trajectories by ob responses in secondary auditory cortex. Journal of Neuroscience 40. 10.1523/JNEUROSCI.2665-19.2020.

28. Liu, H.X., Lopatina, O., Higashida, C., Fujimoto, H., Akther, S., Inzhutova, A., Liang, M., Zhong, J., Tsuji, T., Yoshihara, T., et al. (2013). Displays of paternal mouse pup retrieval following communicative interaction with maternal mates. Nat Commun 4. 10.1038/ncomms2336.

29. Tachikawa, K.S., Yoshihara, Y., and Kuroda, K.O. (2013). Behavioral transition from attack to parenting in male mice: A crucial role of the vomeronasal system. Ann Intern Med 158. 10.1523/JNEUROSCI.2364-12.2013.

30. Vom Saal, F.S. (1985). Time-contingent change in infanticide and parental behavior induced by ejaculation in male mice. Physiol Behav 34, 7–15. 10.1016/0031-9384(85)90069-1.

31. Vom Saal, F.S., and Howard, L.S. (1982). The regulation of infanticide and parental behavior: Implications for reproductive success in male mice. Science (1979) 215. 10.1126/science.7058349.

32. Froemke, R.C., and Young, L.J. (2021). Oxytocin, Neural Plasticity, and Social Behavior. Annu Rev Neurosci 44, 359–381. 10.1146/annurev-neuro-102320-102847.

33. Carcea, I., Caraballo, N.L., Marlin, B.J., Ooyama, R., Riceberg, J.S., Mendoza Navarro, J.M., Opendak, M., Diaz, V.E., Schuster, L., Alvarado Torres, M.I., et al. (2021). Oxytocin neurons enable social transmission of maternal behaviour. Nature 596. 10.1038/s41586-021-03814-7.

34. Nagasawa, M., Okabe, S., Mogi, K., and Kikusui, T. (2012). Oxytocin and mutual communication in mother-infant bonding. Front Hum Neurosci 6. 10.3389/fnhum.2012.00031.

35. Scott, N., Prigge, M., Yizhar, O., and Kimchi, T. (2015). A sexually dimorphic hypothalamic circuit controls maternal care and oxytocin secretion. Nature 525. 10.1038/nature15378.

36. Insel, T.R. (1990). Regional Changes in Brain Oxytocin Receptors Post-Partum: Time-Course and Relationship to Maternal Behaviour. J Neuroendocrinol 2. 10.1111/j.1365-2826.1990.tb00445.x.

37. Mitre, M., Kranz, T.M., Marlin, B.J., Schiavo, J.K., Erdjument-Bromage, H., Zhang, X., Minder, J., Neubert, T.A., Hackett, T.A., Chao, M. V., et al. (2017). Sex-Specific Diberences in Oxytocin Receptor Expression and Function for Parental Behavior. Gend Genome 1. 10.1089/gg.2017.0017.

38. Ehret, G., and Buckenmaier, J. (1994). Estrogen-receptor occurrence in the female mouse brain: Ebects of maternal experience, ovariectomy, estrogen and anosmia. J Physiol Paris 88. 10.1016/0928-4257(94)90012-4.

39. Koch, M., and Ehret, G. (1989). Immunocytochemical localization and quantitation of estrogen-binding cells in the male and female (virgin, pregnant, lactating) mouse brain. Brain Res 489. 10.1016/0006-8993(89)90012-7.

40. Doerr, H.K., Siegel, H.I., and Rosenblatt, J.S. (1981). Ebects of progesterone withdrawal and estrogen on maternal behavior in nulliparous rats. Behav Neural Biol 32, 35–44. 10.1016/S0163-1047(81)90242-9.

41. Banerjee, S.B., and Liu, R.C. (2013). Storing maternal memories: Hypothesizing an interaction of experience and estrogen on sensory cortical plasticity to learn infant cues. Preprint, 10.1016/j.yfrne.2013.07.008 10.1016/j.yfrne.2013.07.008.

42. Smiley, K.O., Brown, R.S.E., and Grattan, D.R. (2022). Prolactin action is necessary for parental behavior in male mice. The Journal of Neuroscience, JN-RM-0558–22. 10.1523/JNEUROSCI.0558-22.2022.

43. Stagkourakis, S., Smiley, K.O., Williams, P., Kakadellis, S., Ziegler, K., Bakker, J., Brown, R.S.E., Harkany, T., Grattan, D.R., and Broberger, C. (2020). A Neuro-hormonal Circuit for Paternal Behavior Controlled by a Hypothalamic Network Oscillation. Cell 182, 960–975.e15. 10.1016/j.cell.2020.07.007.

44. Liang, M., Zhong, J., Liu, H.X., Lopatina, O., Nakada, R., Yamauchi, A.M., and Higashida, H. (2014). Pairmate-dependent pup retrieval as parental behavior in male mice. Front Neurosci. 10.3389/fnins.2014.00186.

45. Insel, T.R., Preston, S., and Winslow, J.T. (1995). Mating in the monogamous male: Behavioral consequences. Physiol Behav 57, 615– 627. 10.1016/0031-9384(94)00362-9.

46. Batty, J. (1978). Acute changes in plasma testosterone levels and their relation to measures of sexual behaviour in the male house mouse (Mus musculus). Anim Behav 26, 349–357. 10.1016/0003-3472(78)90053-2.

47. Li, M.-J., and Xu, X.-H. (2023). Neural representation of sexual satiety in mice. Zool Res 44, 522–524. 10.24272/j.issn.2095-8137.2023.110.

48. Zilkha, N., Scott, N., and Kimchi, T. (2017). Sexual Dimorphism of Parental Care: From Genes to Behavior. Annu Rev Neurosci 40. 10.1146/annurev-neuro-072116-031447.

49. Vinograd, A., Fuchs-Shlomai, Y., Stern, M., Mukherjee, D., Gao, Y., Citri, A., Davison, I., and Mizrahi, A. (2017). Functional Plasticity of Odor Representations during Motherhood. Cell Rep 21. 10.1016/j.celrep.2017.09.038.

50. Kopel, H., Schechtman, E., Groysman, M., and Mizrahi, A. (2012). Enhanced synaptic integration of adult-born neurons in the olfactory bulb of lactating mothers. Journal of Neuroscience 32. 10.1523/JNEUROSCI.6354-11.2012.

51. Lecca, S., Congiu, M., Royon, L., Restivo, L., Girard, B., Mazaré, N., Bellone, C., Telley, L., and Mameli, M. (2023). A neural substrate for negative abect dictates female parental behavior. Neuron 111, 1094–1103.e8. 10.1016/j.neuron.2023.01.003.

52. Schiavo, J.K., Valtcheva, S., Bair-Marshall, C.J., Song, S.C., Martin, K.A., and Froemke, R.C. (2020). Innate and plastic mechanisms for maternal behaviour in auditory cortex. Nature 587. 10.1038/s41586-020-2807-6.

53. Horrell, N.D., Acosta, M.C., and Saltzman, W. (2021). Plasticity of the paternal brain: Ebects of fatherhood on neural structure and function. Dev Psychobiol 63, 1499–1520. 10.1002/dev.22097.

54. Feldman, R., Braun, K., and Champagne, F.A. (2019). The neural mechanisms and consequences of paternal caregiving. Nat Rev Neurosci 20, 205–224. 10.1038/s41583-019-0124-6.

55. Ehret, G., and Bernecker, C. (1986). Low-frequency sound communication by mouse pups (Mus musculus): wriggling calls release maternal behaviour. Anim Behav 34, 821–830. 10.1016/S0003-3472(86)80067-7.

56. Bridges, R.S. (2015). Neuroendocrine regulation of maternal behavior. Front Neuroendocrinol 36, 178–196. 10.1016/j.yfrne.2014.11.007.

57. Ammari, R., Monaca, F., Cao, M., Nassar, E., Wai, P., Del Grosso, N.A., Lee, M., Borak, N., Schneider-Luftman, D., and Kohl, J. (2023). Hormone-mediated neural remodeling orchestrates parenting onset during pregnancy. Science (1979) 382, 76–81. 10.1126/science.adi0576.

58. Gustafson, P., Ladyman, S.R., McFadden, S., Larsen, C., Khant Aung, Z., Brown, R.S.E., Bunn, S.J., and Grattan, D.R. (2020). Prolactin receptor-mediated activation of pSTAT5 in the pregnant mouse brain. J Neuroendocrinol 32. 10.1111/jne.12901.

59. Ali, S., and Ali, S. (1998). Prolactin Receptor Regulates Stat5 Tyrosine Phosphorylation and Nuclear Translocation by Two Separate Pathways. Journal of Biological Chemistry 273, 7709–7716. 10.1074/jbc.273.13.7709.

60. Brown, R.S.E., Aoki, M., Ladyman, S.R., Phillipps, H.R., Wyatt, A., Boehm, U., and Grattan, D.R. (2017). Prolactin action in the medial preoptic area is necessary for postpartum maternal nursing behavior. Proceedings of the National Academy of Sciences 114, 10779–10784. 10.1073/pnas.1708025114.

61. de Jong, T.R., Chauke, M., Harris, B.N., and Saltzman, W. (2009). From here to paternity: Neural correlates of the onset of paternal behavior in California mice (Peromyscus californicus). Horm Behav 56, 220–231. 10.1016/j.yhbeh.2009.05.001.

62. Wang, Z.X., Liu, Y., Young, L.J., and Insel, T.R. (2000). Hypothalamic vasopressin gene expression increases in both males and females postpartum in a biparental rodent. J Neuroendocrinol 12. 10.1046/j.1365-2826.2000.00435.x.

63. Moreno, A., Rajagopalan, S., Tucker, M.J., Lunsford, P., and Liu, R.C. (2023). Hearing Vocalizations during First Social Experience with Pups Increase Bdnf Transcription in Mouse Auditory Cortex. Neural Plast 2023, 1–13. 10.1155/2023/5225952.

64. Winters, C., Gorssen, W., Ossorio-Salazar, V.A., Nilsson, S., Golden, S., and D’Hooge, R. (2022). Automated procedure to assess pup retrieval in laboratory mice. Sci Rep 12, 1663. 10.1038/s41598-022-05641-w.

65. Stern, J.M., and Mackinnon, D.A. (1976). Postpartum, hormonal, and nonhormonal induction of maternal behavior in rats: Ebects on T-maze retrieval of pups. Horm Behav 7. 10.1016/0018-506X(76)90036-2.

66. Dulac, C., O’Connell, L.A., and Wu, Z. (2014). Neural control of maternal and paternal behaviors. Science (1979) 345, 765–770. 10.1126/science.1253291.

67. Kohl, J., Autry, A.E., and Dulac, C. (2017). The neurobiology of parenting: A neural circuit perspective. BioEssays 39, 1–11. 10.1002/bies.201600159.

68. Saltzman, W., and Ziegler, T.E. (2014). Functional significance of hormonal changes in mammalian fathers. Preprint, 10.1111/jne.12176 10.1111/jne.12176.

69. Bales, K.L., and Saltzman, W. (2016). Fathering in rodents: Neurobiological substrates and consequences for obspring. Horm Behav 77. 10.1016/j.yhbeh.2015.05.021.

70. Natan, R.G., Rao, W., and Geben, M.N. (2017). Cortical Interneurons Diberentially Shape Frequency Tuning following Adaptation. Cell Rep 21, 878–890. 10.1016/j.celrep.2017.10.012.

71. Lakunina, A.A., Nardoci, M.B., Ahmadian, Y., and Jaramillo, S. (2020). Somatostatin-Expressing Interneurons in the Auditory Cortex Mediate Sustained Suppression by Spectral Surround. The Journal of Neuroscience 40, 3564–3575. 10.1523/JNEUROSCI.1735-19.2020.

72. Phillips, E.A.K., Schreiner, C.E., and Hasenstaub, A.R. (2017). Cortical Interneurons Diberentially Regulate the Ebects of Acoustic Context. Cell Rep 20, 771–778. 10.1016/j.celrep.2017.07.001.

73. Zhang, L.I., Tan, A.Y.Y., Schreiner, C.E., and Merzenich, M.M. (2003). Topography and synaptic shaping of direction selectivity in primary auditory cortex. Nature 424, 201–205. 10.1038/nature01796.

74. Ye, C., Poo, M., Dan, Y., and Zhang, X. (2010). Synaptic Mechanisms of Direction Selectivity in Primary Auditory Cortex. The Journal of Neuroscience 30, 1861–1868. 10.1523/JNEUROSCI.3088-09.2010.

75. Tasaka, G., Hagihara, M., Irie, S., Kobayashi, H., Inada, K., Kobayashi, K., Kato, S., Kobayashi, K., and Miyamichi, K. (2025). Orbitofrontal cortex influences dopamine dynamics associated with alloparental behavioral acquisition in female mice. Sci Adv 11. 10.1126/sciadv.adr4620.

76. Gubernick, D.J., and Nelson, R.J. (1989). Prolactin and paternal behavior in the biparental California mouse, Peromyscus californicus. Horm Behav 23, 203–210. 10.1016/0018-506X(89)90061-5.

77. Buonfiglio, D.C., Ramos-Lobo, A.M., Silveira, M.A., Furigo, I.C., Hennighausen, L., Frazão, R., and Donato, J. (2015). Neuronal STAT5 signaling is required for maintaining lactation but not for postpartum maternal behaviors in mice. Horm Behav 71, 60–68. 10.1016/j.yhbeh.2015.04.004.

78. Anderson, G.M., Beijer, P., Bang, A.S., Fenwick, M.A., Bunn, S.J., and Grattan, D.R. (2006). Suppression of Prolactin-Induced Signal Transducer and Activator of Transcription 5b Signaling and Induction of Suppressors of Cytokine Signaling Messenger Ribonucleic Acid in the Hypothalamic Arcuate Nucleus of the Rat during Late Pregnancy and Lactation. Endocrinology 147, 4996–5005. 10.1210/en.2005-0755.

79. Li, X.-Y., and Toyoda, H. (2015). Role of leak potassium channels in pain signaling. Brain Res Bull 119, 73–79. 10.1016/j.brainresbull.2015.08.007.

80. Bean, B.P. (2007). The action potential in mammalian central neurons. Nat Rev Neurosci 8, 451–465. 10.1038/nrn2148.

81. Lyons, D.J., Hellysaz, A., and Broberger, C. (2012). Prolactin Regulates Tuberoinfundibular Dopamine Neuron Discharge Pattern: Novel Feedback Control Mechanisms in the Lactotrophic Axis. Journal of Neuroscience 32, 8074–8083. 10.1523/JNEUROSCI.0129-12.2012.

82. Kamesh, A., Black, E.A.E., and Ferguson, A. V. (2018). The subfornical organ: A novel site for prolactin action. J Neuroendocrinol 30. 10.1111/jne.12613.

83. Goldey, G.J., Roumis, D.K., Glickfeld, L.L., Kerlin, A.M., Reid, R.C., Bonin, V., Schafer, D.P., and Andermann, M.L. (2014). Removable cranial windows for long-term imaging in awake mice. Nat Protoc 9, 2515–2538. 10.1038/nprot.2014.165.

84. Flickinger, D., Iyer, V., Huber, D., O’Connor, D., Peron, S., Clack, N., Chandrashekar, J., and Svoboda, K. (2010). MIMMS: a modular, open design microscopy platform for in vivo imaging of neural tissues. In Soc Neurosci Abstr.

85. Pologruto, T.A., Sabatini, B.L., and Svoboda, K. (2003). ScanImage: Flexible software for operating laser scanning microscopes. Biomed Eng Online 2, 13. 10.1186/1475-925X-2-13.

