## Supplementary figures for "Prolactin Shapes Cortical Plasticity in Fathers"

### Extended data Figure 1

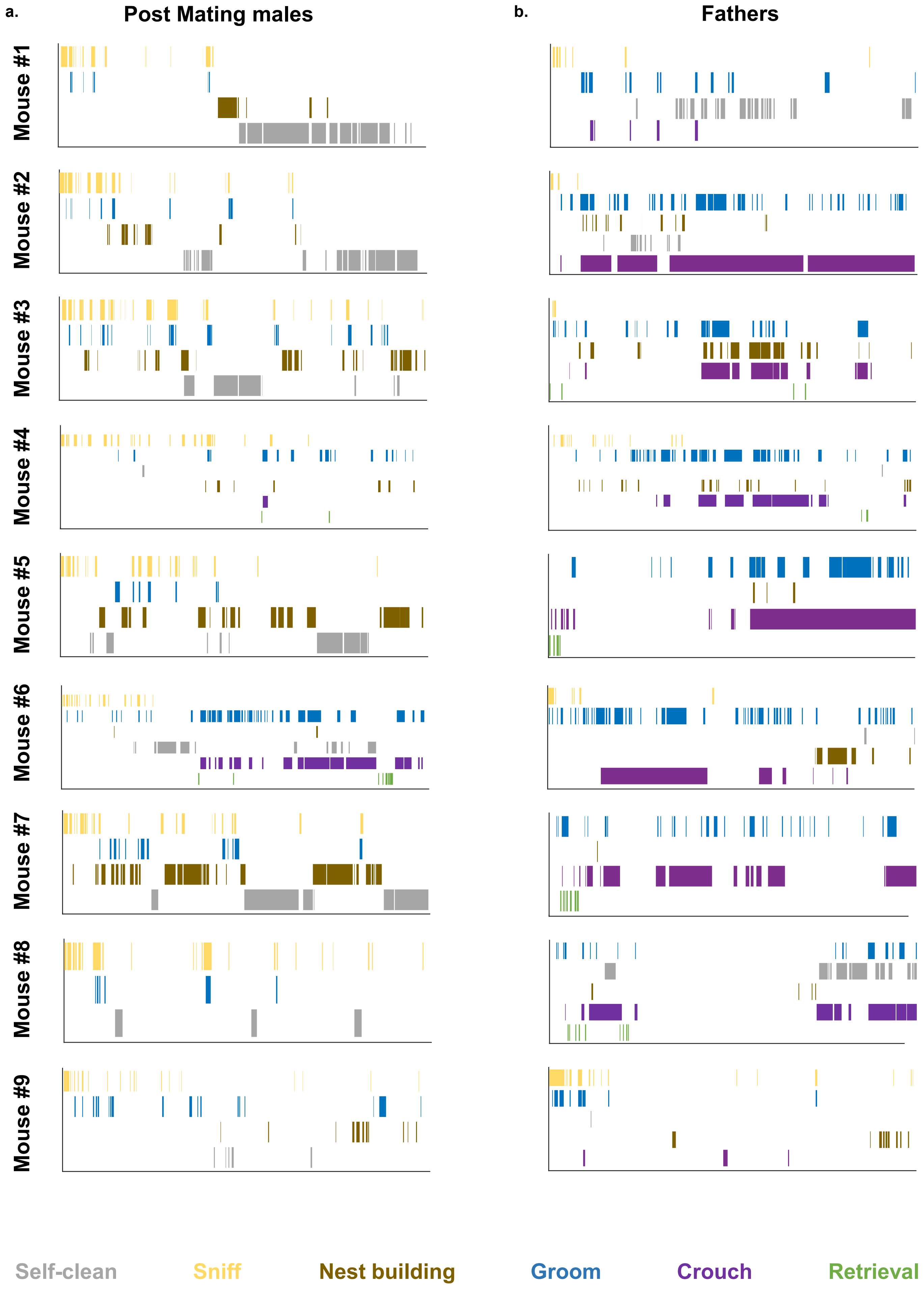

### Extended data Figure 2

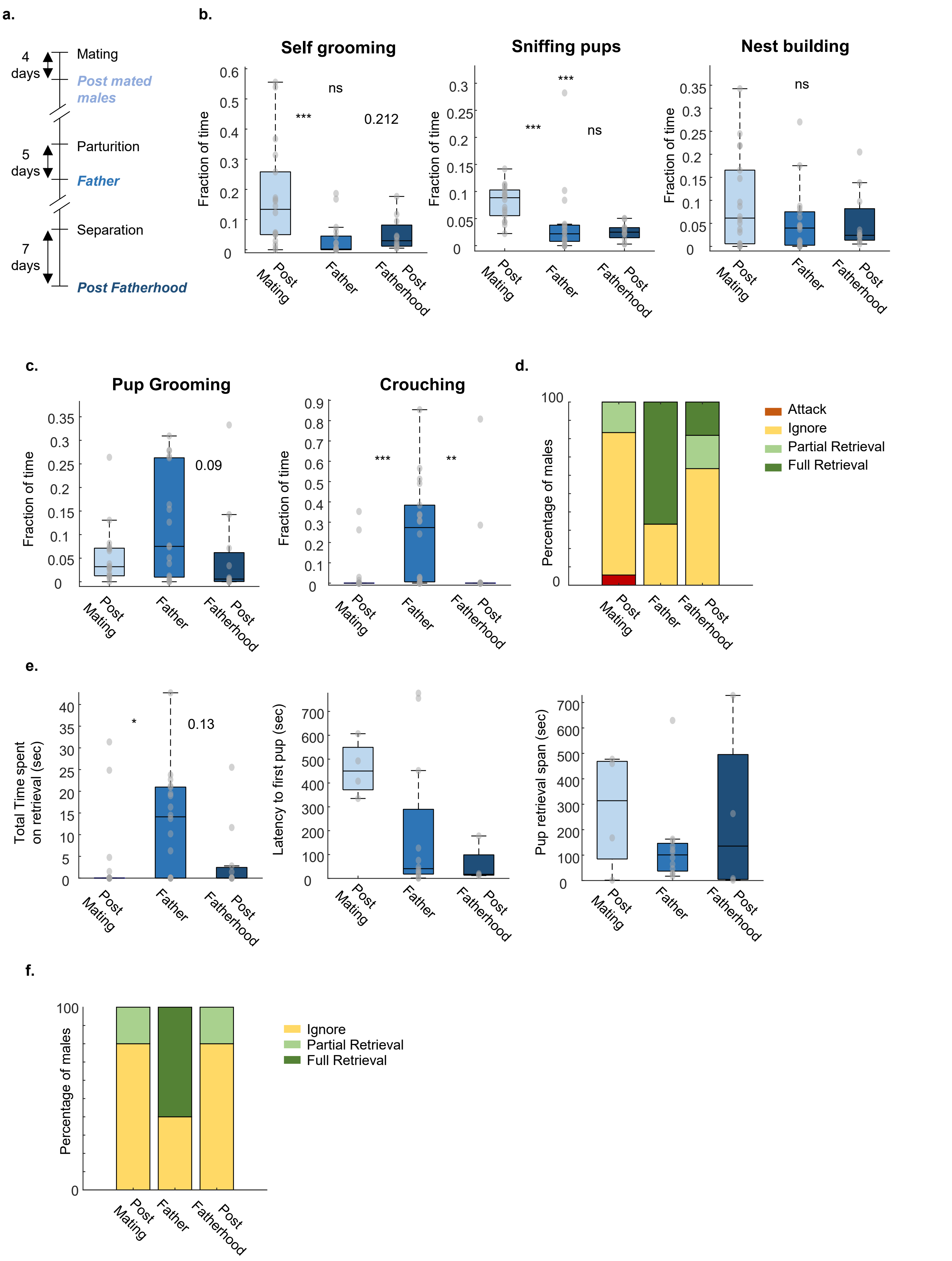

### Extended data Figure 3

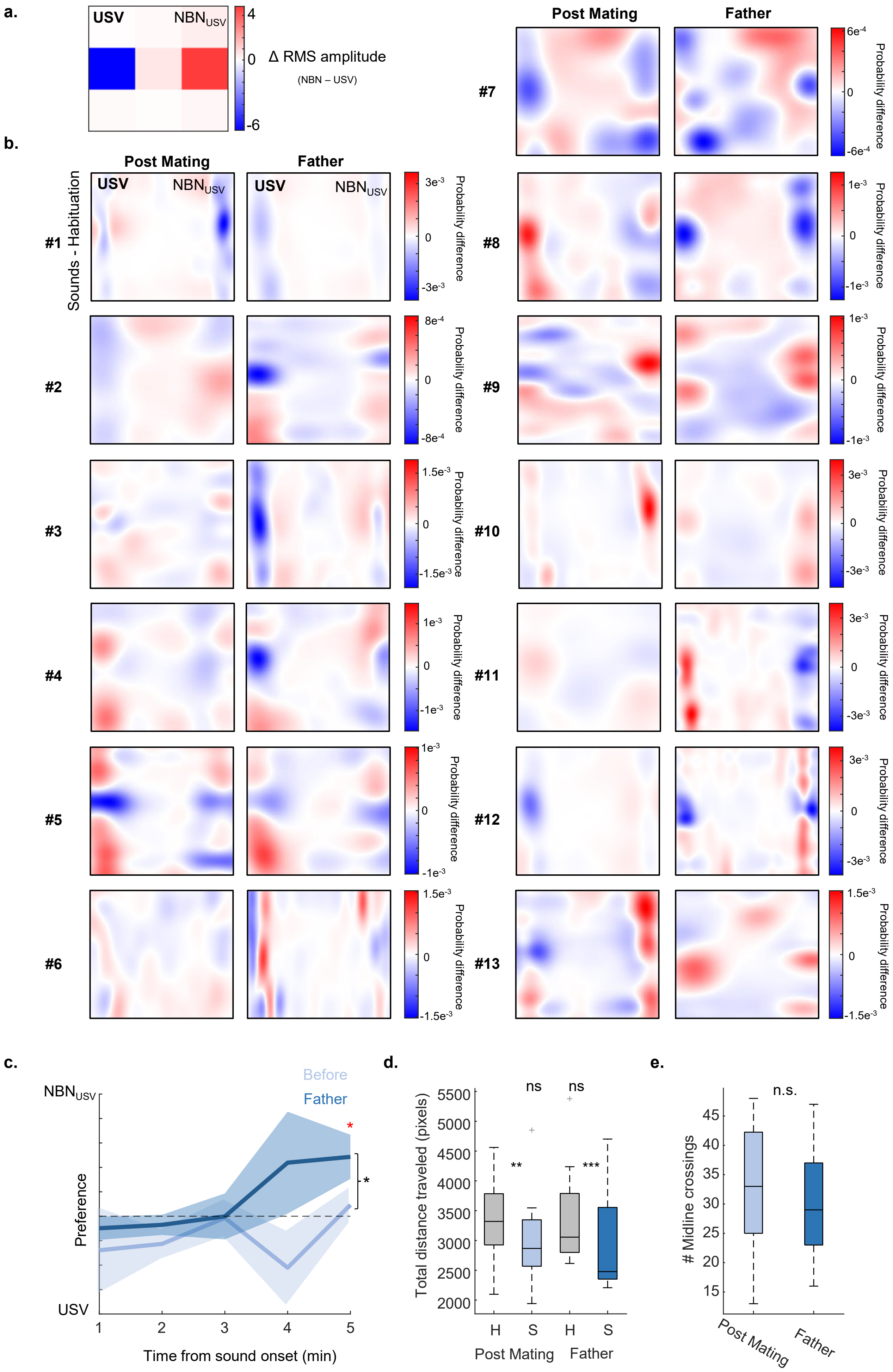

### Extended data Figure 4

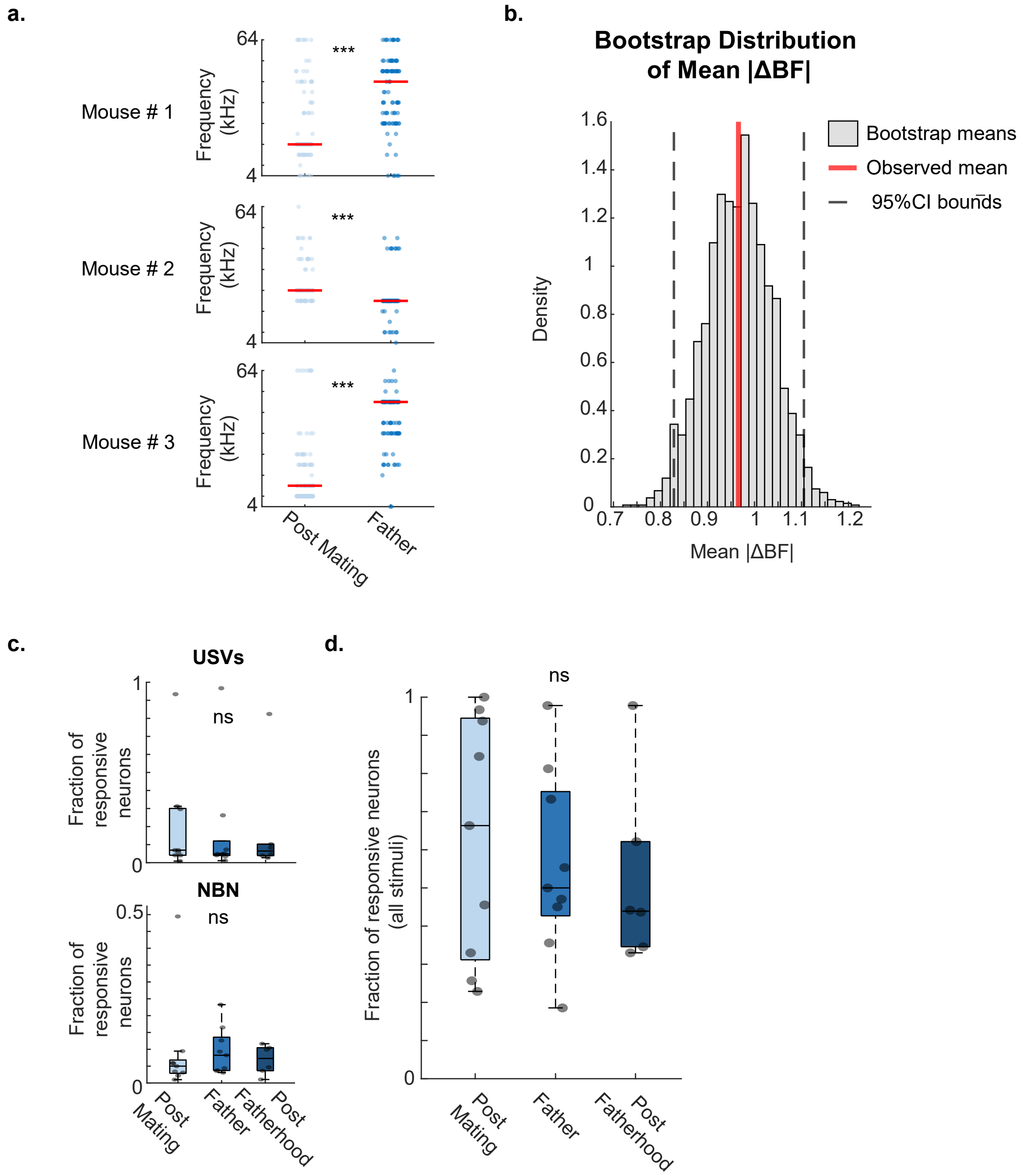

### Extended data Figure 5

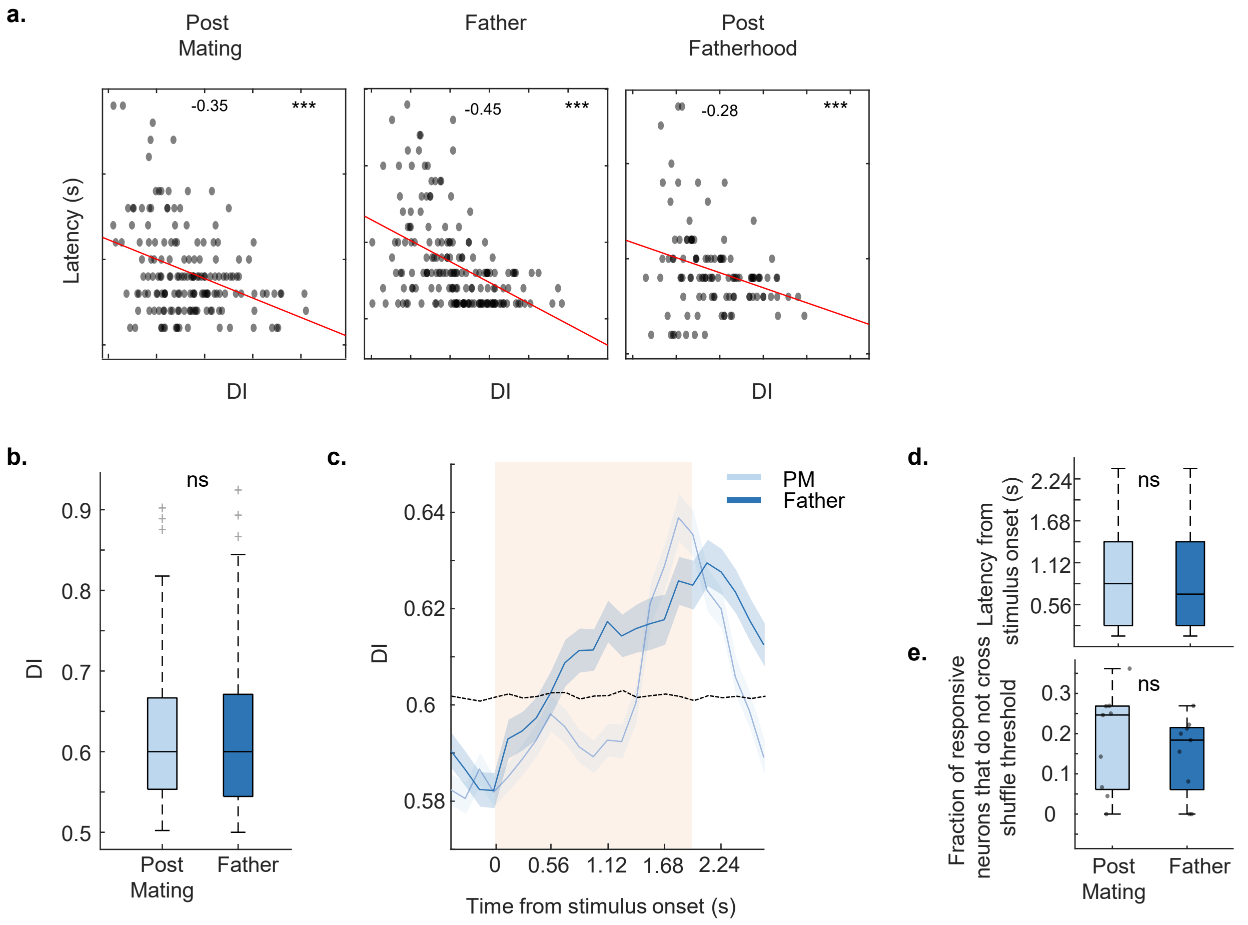

### Extended data Figure 6

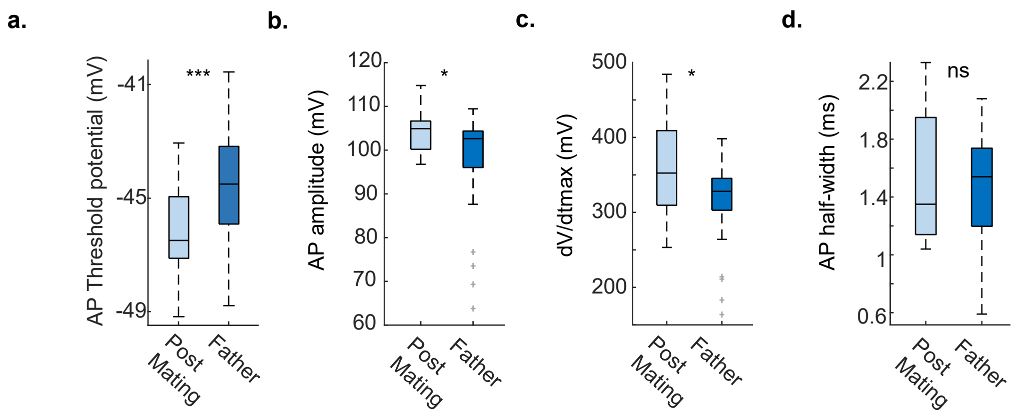

### Extended data Figure 7

a.

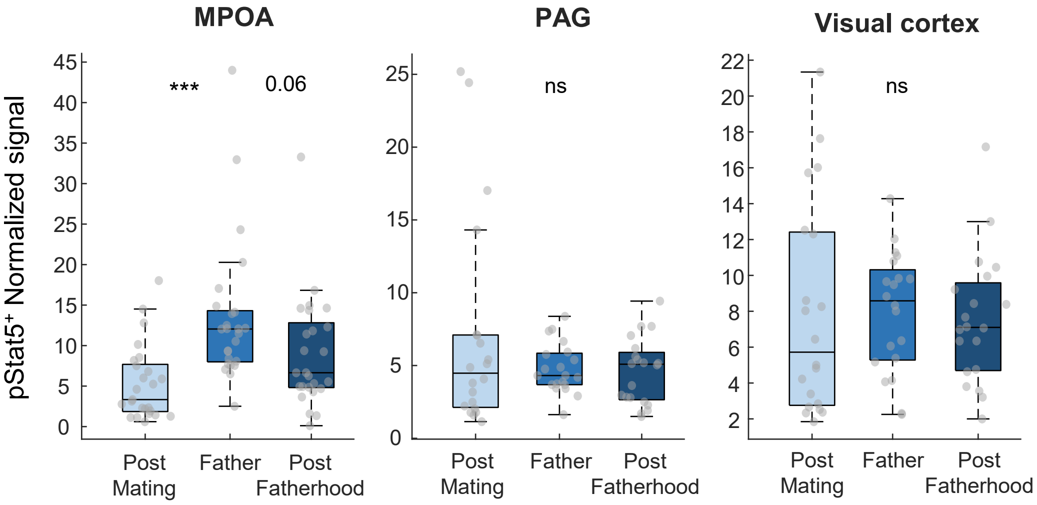

### Extended data Figure 8

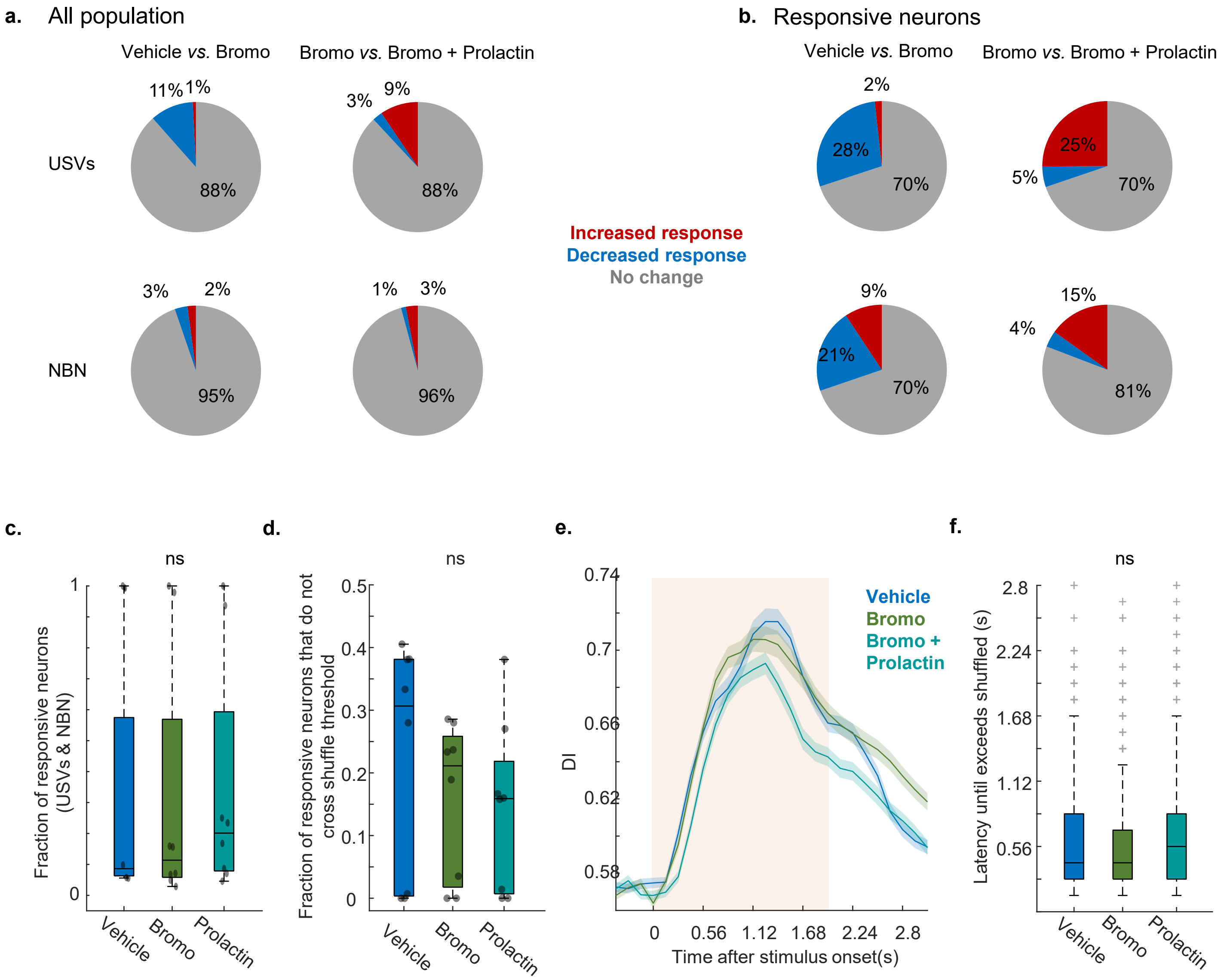
